# A recurrent vision transformer shows signatures of primate visual attention

**DOI:** 10.1101/2024.11.09.622721

**Authors:** Jonathan Morgan, Badr Albanna, James P. Herman

## Abstract

We present a Recurrent Vision Transformer (Recurrent ViT) that integrates a capacity-limited spatial memory module with self-attention to emulate primate-like visual attention. Trained via reinforcement learning on a spatially cued orientation-change detection task, our model exhibits hallmark behavioral signatures of primate attention—including improved detection accuracy and faster reaction times for cued stimuli that scale with cue validity. Analysis of its self-attention maps reveals rich temporal dynamics: spatial biases induced by cues are maintained during blank intervals and reactivated prior to anticipated stimulus changes, mirroring the top–down modulation observed in primate studies. Moreover, targeted manipulations of internal attention weights yield performance changes analogous to those produced by microstimulation in attentional control regions such as the frontal eye fields and superior colliculus. These findings demonstrate that embedding recurrent, memory-driven mechanisms within transformer architectures may provide a computational framework for linking artificial and biological attention

## 1 Introduction

Visual attention is a foundational cognitive function that enhances perceptual judgments and speeds reaction times for attended stimuli [1–5]. Neural correlates include heightened spiking activity and decreased spike-count correlations for attended stimuli [6–10] Classic paradigms use spatial cues to direct attention [11], where a delay separates cue and stimulus; the cue’s location must therefore be maintained in visual working memory (VWM). This requirement reinforces strong links between attention and VWM [12–15], as working memory contents guide attention and vice versa [16–22]. Stored locations can be updated [23–25], and attention can prioritize locations for working memory encoding [12, 26,27] or reorient to memorized locations to strengthen representations [28–31], thereby improving recall [22].

Transformers [32–34] have achieved remarkable success in both language and vision, using self-attention mechanisms reminiscent of how biological systems might allocate attentional resources [35–37]. However, whether their form of “attention” aligns with human attention remains debated. Farxivor instance, transformers can show human-like patterns in text-based tasks [38], but in vision tasks they often emphasize low-level grouping rather than goal-driven selection [39]. Unsupervised training regimes such as DINO can yield attention maps resembling human gaze [40], yet these models typically observe entire sequences of input frames at once, bypassing the limitations of human perceptual or working memory. When continuous access to past stimuli is granted, the burden of selective encoding and storage is diminished; thus, claims of biological plausibility become tenuous if critical constraints like limited capacity [41–44] and dynamic internal states are absent [45–47].

Because standard vision transformers that process only a single current image cannot match basic primate attentional capabilities, one must introduce a top-down guide into their self-attention architecture. To address this, we propose a ViT variant that includes a spatial memory module feeding back into self-attention, integrating ideas from recurrent neural networks [48, 49]. We train it on a cued orientation-change detection task [11,50] in a reinforcement-learning framework. The model exhibits improved performance and faster responses for cued stimuli, paralleling attentional benefits observed in primates [1–5]. Moreover, targeted manipulations of the model’s attention weights yield performance changes reminiscent of those seen following causal manipulations in frontal eye fields [51] and superior colliculus [52,53]. These findings highlight that incorporating spatial memory and feedback into vision transformers can recover core signatures of primate attention, suggesting a promising path for reconciling transformer-based architectures with biological principles of attentional control.

## 2 Task and model

### 2.1 Cued orientation change detection task environment

We trained our model on an orientation change detection task in which the agent must report if a change has occurred during each trial Figure 1. Each trial comprised 7 time steps. At each time step, a 50×50 grayscale image was shown to the agent. At *t* = 0, the trial began with a black image. At *t* = 1, a spatial cue was displayed. The cue appeared at stimulus position 1 (Cue *S*_1_) on one half of the trials and at position 4 (Cue *S*_4_) on the other half. At *t* = 2, a black screen was again displayed. At *t* = 3, stimulus onset occurred, displaying four “Gabor” stimuli in randomly chosen orientations at fixed positions: top left (*S*_1_), bottom left (*S*_2_), top right (*S*_3_), and bottom right (*S*_4_). At *t* = 4, the stimuli remained unchanged with the exception of orientation “noise” added in each time step of stimulus presentation to control task difficulty (see Methods). If the trial was a no-change trial, stimuli remain unchanged from *t* = 4 to *t* = 6. In a change trial, at *t* = 5 the orientation of one of the four stimuli changed by Δ degrees (where Δ varied from trial to trial). The orientations then remained unchanged from *t* = 5 to *t* = 6. Half of all trials were “change trials” and and the other half were “no-change trials” (balanced across cue presentation positions).

**Figure 1.**
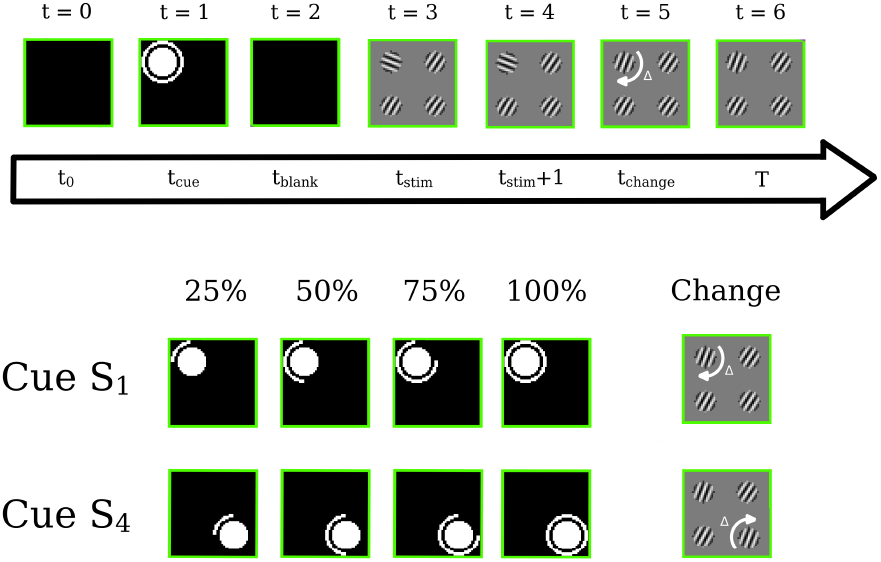
Each trial in the task comprises seven time steps. In each time step, a 50 × 50 grayscale image is input to the model. Black images are shown at *t* = 0 and *t* = 2. The cue is shown at *t* = 1 and can either be at *S*_1_ (top left) or *S*_4_ (bottom right). The cue can take four configurations, where the ring around the center disk changes, indicating the probability (25%, 50%, 75%, or 100% chance) that the change will appear at the cued location if the trial is a change trial.

**Figure 2.**
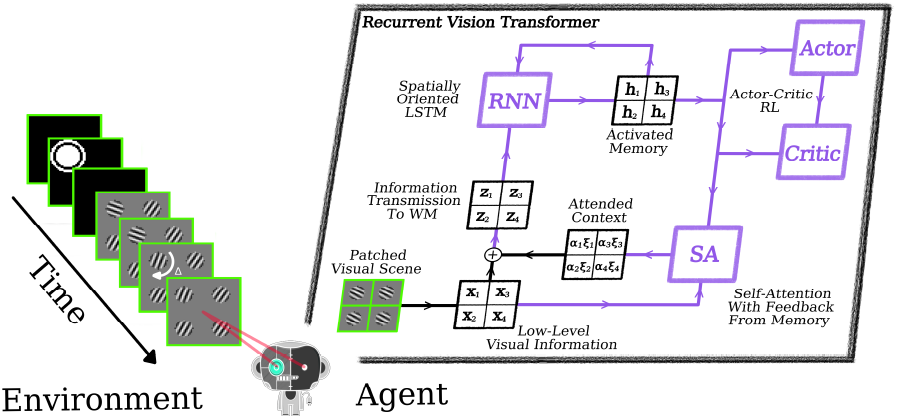
This schematic illustrates how the agent processes visual inputs and updates its internal state using a Recurrent Vision Transformer (Recurrent ViT). At each timestep, the environment provides a single image, which is split into four patches (one for each visual stimulus) and then passed through a pre-processing stage (see Methods). The resulting low-level visual features, 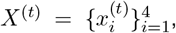, are combined with previously activated memory, 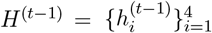, within a self-attention mechanism, producing spatio-temporal context vectors (*α*_*i*_*ξ*_*i*_) that encode essential information about the current scene. These context vectors are added to the low-level features and processed together to yield a new transmission, 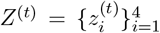. The agent then updates its memory, 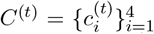,using both *Z*^(*t*)^ and the previously activated memory *H*^(*t*−1)^. A subset of the updated memory is re-activated, *H*^(*t*)^, and fed back into both the self-attention mechanism and the actor and critic networks. The actor network uses this activated memory to determine the action that maximizes future rewards, while the critic network evaluates pairs of activated memory and actions to estimate upcoming cumulative rewards. Purple lines indicate learnable projections (synaptic weights) updated through reward feedback.

Visually distinct cues indicated different levels of “cue validity”: the probability of an orientation-change event occurring at the cued position. Cue validity levels were 25%, 50%, 75%, or 100%. For example, the 50% valid cue presented at *S*_1_ meant that, if this was a change trial, there was a 50% probability that the orientation change would occur at position *S*_1_. Cue validity was depicted visually by a white arc that subtended 25%, 50%, 75% or 100% of a central disc’s circumference (Figure 1). We use the term “cue validity” to align with the convention established in the human and NHP psychophysics literature [11, 54–56].

Much like NHPs trained in visual attention tasks, we train our model in an RL setting. At each time step the agent could either choose to wait (*a*^(*t*)^ = ‘wait’) or declare a change (*a*^(*t*)^ = ‘declare change’). For *t <* 6, waiting resulted in no reward (*r* = 0), and the trial advanced to the next time step. At *t* = 6, waiting rewarded the agent *r* = 1 in a no-change trial and *r* = 0 in a change trial. Declaring a change always ended the trial. When the agent correctly declared a change at *t*≥ 5, it received a reward *r* = 1. In all other cases, declaring a change resulted in *r* = 0. Before describing the model’s behavior, we first briefly describe model components and their interconnections to facilitate interpretation of the model’s attention dynamics.

### 3 Model

Our Recurrent ViT comprises two modules: the self-attention (SA) module and the patch-based long-short-term-memory (LSTM) module. We train the model in an actor-critic RL framework. The environment generates the current visual scene and the agent converts this scene to a visual representation, *X*^(*t*)^, through low-level convolutional operations (see supplement). The low-level processing, results in 4 visual patches: 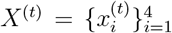,and this 4-patch structure is retained throughout the attention and working memory modules (number of patches is a hyperparameter). The primary benefit of our choice to structure the visual patches around stimulus positions is the interpretability it affords to the model’s “attention map”. Specifically, it allows the attention map to be visualized as a 4×4 array in which we can interpret as the bias, 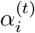,assigned to an internal representation 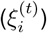 associated with stimulus *S*_*i*_. An important distinction here is that 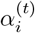 does not just describe the the attention on the current visual patch 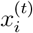.Instead it describes the attention on an internal representation, 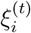 that consists of the immediate visual information in 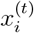 and information derived from activated memory describing relevant past temporal and spatial context, 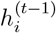.

The patch-based LSTM module receives a transmission, 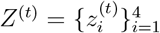, that contains visual information derived from the immediate visual scene in addition to spatial and temporal context derived from the selfattention mechanism. This information is utilized to update the internal states of the LSTM, 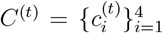 (see Methods). Activated memory, 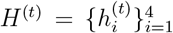, is derived from these internal states and sent: (1) recurrently back into the LSTM; (2) to the self-attention module; (3) to the actor and critic networks. This allows attention to be allocated both on the basis of the next visual inputs, *X*^(*t*+1)^, as in traditional transformer architectures [32, 33], and on the basis of memory (*C*^(*t*)^). A full, detailed description of the model can be found in the Methods section.

## 4 Results

### 4.1 A recurrent ViT exhibits behavior signature of visual attention

Our model exhibited orderly “psychometric” and “chronometric” functions with characteristic sigmoidal shapes commonly observed in human and NHP experiments (Figure 3A,D). Larger orientation-change Δ values were associated with increased hit rates and decreased reaction times, qualitatively comparable to those seen in countless human and NHP psychophysics experiments. For cued orientation changes, higher cue validity improved correct response rate (Figure 3A) and sped reaction time modestly (Figure 3D). This pattern mirrors experimental findings that attentional benefits in biological systems are most pronounced when discriminating subtle changes. No effects of cue validity were observed on fitted psychometric function slope, guess rate, or lapse rate, indicating that, like spatial attention in biological visual systems, our model’s attention mechanism produced primarily additive effects on perceptual sensitivity rather than changing the shape of the psychometric function (Supplement) [57–59].

**Figure 3.**
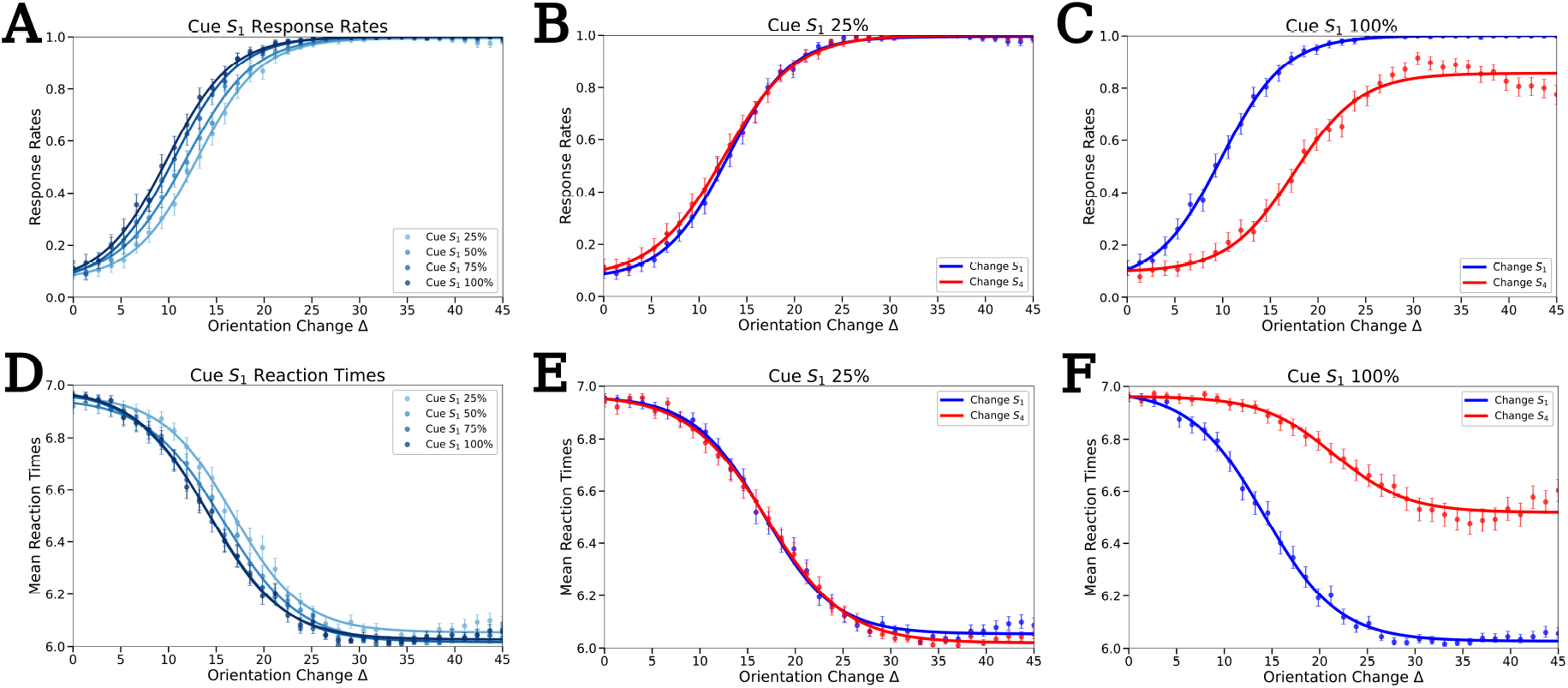
**A–F** shows the response rates (**A–C**) and reaction times (**D–F**) of our agent over varying cue validities with respect to the *S*_1_ location and either a change on *S*_1_ or a change on *S*_4_ positions. Each data point was 500 trials where Δ specifies the magnitude of the orientation change. The response-rate was computed as *n*_dc_*/n*_trials_, where *n*_dc_ is the total number of trials in which the agents selected the action *a*^(*t*)^ = “declare change” and *n*_*trials*_ is the total number of trials. The reaction times were computed as 1*/*500 ∑_*i*_ *τ*_*i*_, where *τ*_*i*_ is the time the trial ended, either by the agent declaring a change or waiting through the final timestep. **A** Response rates over each possible cue condition w.r.t. the *S*_1_ position where changes also occurred at the *S*_1_ position. **B** Response rates computed over trials with a cue at the *S*_1_ position comparing changes at the *S*_1_ versus the *S*_4_ locations. **C** Similar to **B** but with a 100 % cue at the *S*_1_ location. **D–F** Same conditions as **A–C** showing the mean reaction times.

Contrasting performance in response to cued orientation changes compared to uncued revealed a clear “cueing effect”, again recapitulating results from human and NHP literature [1,5,60–65]. Using the trained model with fixed weights, we were able to test uncued conditions even in the 100% cue validity case, despite the absence of these trials during training (Figure 3C,F). In that 100% cue validity condition, for example, 10° cued orientation changes were detected in roughly 50% of trials, but uncued changes of the same magnitude were detected in roughly 15% of trials (Figure 3C). The magnitude of this cueing effect varied systematically with cue validity: compared to the 100% validity condition, cue effects were mostly absent in the 25% condition (Figure 3B,C), which is expected - because there are 4 stimulus positions, 25% cue validity indicates equal probability of the orientation change occurring at any stimulus position. Thus, the 25% valid cue should not confer any selective preference for one stimulus over another.

### 4.2 A recurrent ViT deploys attention in an human/NHP-like strategy

In Figure 4A, we present averaged self-attention heatmaps from trials with the cue at location *S*_1_ and and no orientation changes of Δ = 0. Prior to presentation of any visual stimuli (*t* = 0), the model’s attention landscape is flat (Figure 4A). When the spatial cue is presented (*t* = 1), attention is strongly biased 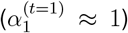 to the cued location. This bias persists during the blank interval (*t* = 2), indicating that the activated memory 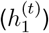 can maintain spatial attention even in the absence of meaningful visual input. At stimulus onset (*t* = 3), attention maps become largely flat again, with a slight off-diagonal bias. Immediately before the time of orientation change (*t* = 4), the model begins to weakly allocate attention to the cued location, with stronger attention bias for higher cue validity. This mirrors attention dynamics observed in humans and animals performing tasks in which sensory events occur at predictable times [66–70]. When the cued orientation change occurs (*t* = 5), we see the integration of stimulus-driven attention and memory-derived attention. Specifically, even with no orientation change occurring, higher cue validity leads to stronger attention bias toward the cued location at the time of the change (*t* = 5). This attention bias lingers into the subsequent time step (*t* = 6), with magnitude also varying with cue validity. These results are consistent with theories of attention control that postulate the integration of multiple sources of attention bias [71–76], and clearly demonstrate how our Recurrent ViT model has learned to allocate attention on the basis of both the predictability offered by the spatial cue and on the basis of the behavioral relevance of features in the input image(s).

**Figure 4.**
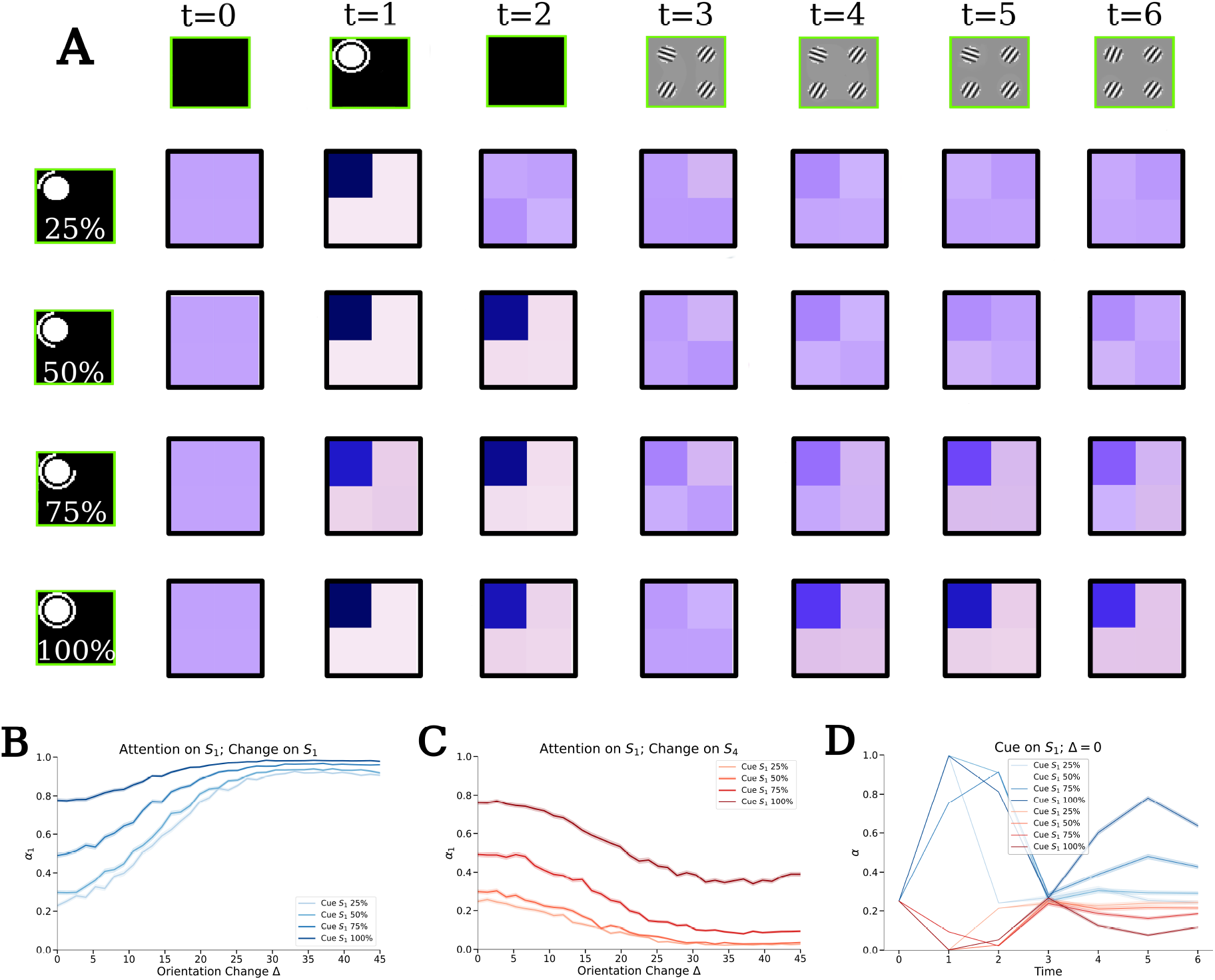
**A** Averaged self-attention maps at each timestep when *S*_1_ is cued (various cue validities) and Δ = 0. Rows (top to bottom) indicate different cue validities (25%, 50%, 75%, 100%), and columns (left to right) show distinct timesteps *t* = 0 6. Darker squares in each 2 × 2 attention map reflect stronger attention. **B–C** Attentional bias 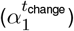 assigned to the *S*_1_ position over a vaying orientation change magnitude Δ and varying cue validity. The bias 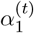 is the top-left value shown qualitatively in the heatmaps from **A. B** Blue coloring represents the varying cue validity, with darker blues being higher validity. **C** Red coloring represents the varying cue validity, with darker red being lower validity. **D** The spatial bias (blue: 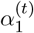 and red: 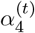)at different time points and over varying cue validity on trials with no orientation change.

Having found that both orientation change and cue validity contribute to attention allocation, we next sought to quantify how these two factors interact. To this end, we computed 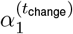 (time of change: *t*_change_ = 5) separately for each cue validity condition on trials in which we cued *S*_1_ and systematically varied the orientation change Δ presented at either *S*_1_ (Figure 4B) or *S*_4_ (Figure 4C). When the change was at the cued location (*S* ), 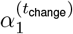 increased sigmoidally with larger Δ values, with the asymptotic value scaling with cue validity (Figure 4B blue coloring). Conversely, when the change occured at the uncued location (*S*_4_ ), larger Δ values decreased 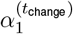as attention was drawn to the change location. The magnitude of this decrease depended systematically on cue validity: With 25% valid cues, large changes at *S*_4_ drove 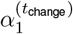 near 0, indicating strong attentional capture by the uncued change. However, with increasing cue validity (Figure 4C red coloring), 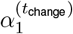 maintained progressively higher values even for large Δ at *S*_4_, demonstrating that strongly predictive cues can partially maintain attention at the cued location in the face of competing visual events.

Finally, in Figure 4D we show the attentional bias on 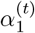 (blue lines) and 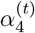 (red lines) for different cue *S*_1_ validities and each timestep in the trial. Each of the trials used to collect this data were trials in which there was not an orientation change. We found that attentional modulation peaked at the time of the cue, with comparable biases independent of the cue validity. For cue validities corresponding to the location of the attentional bias being measured (Cue *S*_1_ and measuring *α*_1_), the bias increases (blue lines at *t* = 1). Due to the finite attentional resource in our model, this is accompanied with a decrease in attentional biases at other locations (red lines). Attentional modulations remain elevated (suppressed) following the cue (*t* = 2) even when only a blank visual scene is shown (not including the 25% condition). At stimulus onset (*t* = 3), the attentional bias returns to roughly uniform distribution, regardless of the cue condition. Then at *t* = 4 and *t* = 5, the attentional bias increases (decreases for *α*_4_) and reach a maximum at *t* = 5 before decreasing at *t* = 6. Furthermore, the higher the cue validities, the sharper this increase. The relative suppression of attentional bias at stimulus onset, the sharp rise in attentional bias with high validity cues following stimulus onset, and the decrease in attentional biases on the last step of the trial resemble experimental results taken from neuronal activity in area V4 of NHPs also solving a change detection task [67].

### 4.3 Manipulating bias affects response rate and reaction times

Experimental studies have shown that microstimulation of brain regions involved in attentional control, such as the frontal eye fields (FEF), lateral intraparietal area (LIP), and superior colliculus (SC), can bias visual processing toward specific retinotopic locations in the visual scene [51–53,77]. For example, microstimulation of the FEF enhances the neural representation of stimuli at corresponding spatial locations, leading to improved detection performance and faster reaction times for stimuli presented at those locations [51]. Similarly, microstimulation of the SC has been shown to modulate both perceptual sensitivity and decision criterion, affecting how stimuli are processed and how responses are generated [53].

In our model, artificially increasing the bias 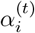 assigned to specific spatio-temporal representation, 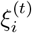,can generate similar effects. This results in the prioritization of information from the visual patch 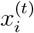 and the activated memory 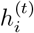.As demonstrated in Figure 5A-C, when we artificially increase the bias in favor of the change location (e.g., setting 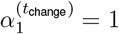), we observe a slight increase in hit rates for detecting the change. This suggests that enhancing the attentional bias toward the relevant location facilitates the processing of the stimulus change, leading to improved detection performance.

**Figure 5.**
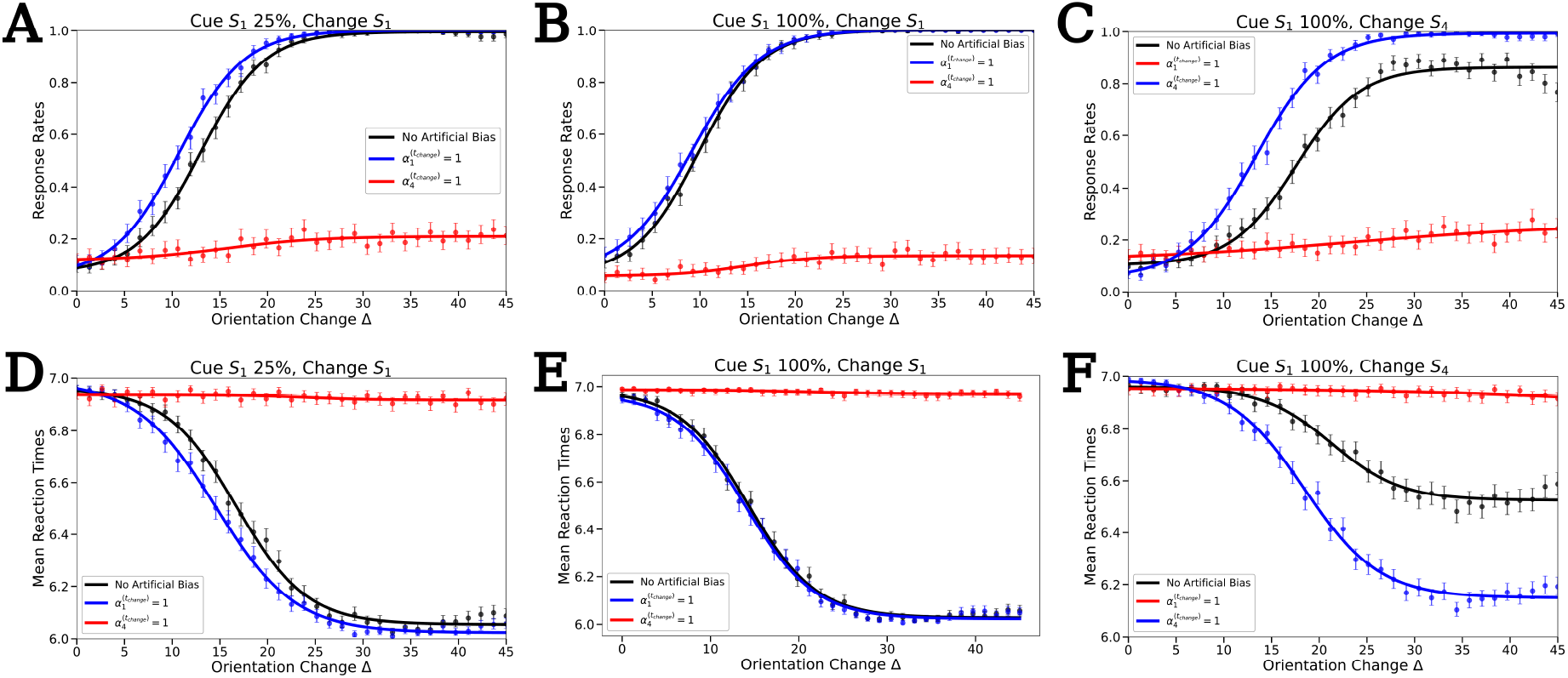
Plots showing the effect of artificially modulating the bias. All data points are the result of an average over 500 trials. Artificial modulation involves inducing a high bias in a single spatial region (increasing the value of one of the patches in the self-attention maps from Figure 4G). In all cases, the bias is induced with respect to the 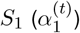 or te 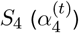 stimulus location. In **A–F**, we plot the response rates and reaction times versus the orientation change Δ. If 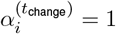, this indicates that the transmission 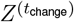 has been completely biased toward 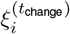.In **A–C** we show the effects of this manipulation on the response rates, and then again for the reaction times in **D–F**.

Conversely, when we increase the bias toward a location different from where the change occurs, we see a significant decrease in response rates (Figure 5A-C). Directing bias away from the relevant stimulus location where a change occurred impairs detection performance, due to the spatio-temporal representation with respect to the change location being absent in the transmission to the working memory module. The inhibitory effect of misallocated attention is further reflected in reaction times (Figure 5D-F), where inducing a high bias at non-change locations results in significant increases in response latency. These findings are consistent with behavioral studies showing that invalid spatial cues or misdirected attention can slow down responses and reduce accuracy [11,54–56].

## 5 Discussion

In this work, we introduced a Recurrent Vision Transformer (Recurrent ViT) equipped with a capacity-limited spatial memory module to perform a cued orientation change-detection task. Our aim was to address the question of whether standard vision transformers—which often rely on either feedforward processing of single frames [33] or post-hoc attention maps [40]— exhibit the kind of top-down, internally guided attentional mechanisms that are fundamental to human and non-human primate (NHP) vision [1,39,72]. . By integrating a recurrent, spatial working memory into the model, we found that our Recurrent ViT recapitulates many hallmark effects of primate visual attention.

### 5.1 Recovering hallmark signatures of primate attention

First, our trained model shows improved performance and faster detection of orientation changes at cued locations, mirroring the well-documented behavioral effects of selective spatial attention [1, 3, 5, 75]. These benefits emerge in situations where high-validity cues bias internal representations toward the cued location, but they taper off or reverse if competing salience signals (e.g., a large orientation change elsewhere) dominate the model’s self-attention. This interplay between cue validity and exogenous salience resonates with empirical observations that attentional allocation reflects both top-down predictions and bottom-up feature-driven signals [71, 72]. In standard feed-forward ViTs, attention is inherently limited to correlational or grouping-based processes [39, 78], whereas our recurrent module explicitly integrates memories of cue identity, location, and temporal context—restoring top-down selectivity typically absent in off-the-shelf architectures.

### 5.2 Relevance to neural mechanisms of attention and working memory

The success of our Recurrent ViT underscores the deep links between attention and working memory reported in neuroscience [12, 14, 15, 24]. Much like the “attentional template” theory, which proposes that memory representations guide attention to relevant features and locations [26,27,59], our model maintains a set of spatial codes over time. These memory states re-enter a self-attention module to bias ongoing visual processing, effectively bridging top-down and bottom-up circuits [51, 53, 79, 80]. Correspondingly, the interplay between memory and perception in our model echoes the reciprocal loops seen in primate frontal, parietal, and subcortical structures, where neural firing maintains spatial priority during blank intervals and facilitates rapid reactivation at anticipated moments of stimulus change [50, 81, 82].

### 5.3 Subcortical and dopaminergic influences

Although our model already uses reward feed-back to guide learning, we have not explicitly integrated dopaminergic-like prediction error signals or examined how reward history might adaptively modulate attentional policies in the superior colliculus and related circuits [83–86]. In biological systems, dopamine critically mediates plasticity, enabling more nuanced shaping of attentional priorities over extended time scales [87–89]. A potential future extension is to incorporate a free-energy principle-inspired, unsupervised component [90–94], which could allow the model to learn latent, generative structure in its environment—paralleling how dopamine modulates not only immediate reward but also uncertainty and exploration in real brains. Such a framework would unify reward-driven reinforcement learning with a broader predictive coding approach, further enhancing the model’s capacity for dynamic, context-sensitive attention.

### 5.4 Constraints, biological plausibility, and interpretability

A critical innovation in our model is the deliberate imposition of a capacity-limited spatial memory, distinguishing it from standard transformers that can re-attend to entire sequences without the bottlenecks characteristic of primate vision [32, 33, 95] and from vision transformers that take entire sequences of images as input [40]. This bottleneck fosters internal competition for representational resources, consistent with models of biased competition that posit a finite “priority map” mediating competition between stimuli and top-down goals [5, 76, 79, 96, 97]. By structuring our LSTM such that each spatial patch is assigned only a single hidden-state slot, our Recurrent ViT is compelled to compress or discard irrelevant information on every timestep. This capacity-limited design parallels well-known psychophysical findings that visual working memory is resource limited [41–44]. In practice, the model learns to devote its memory resources primarily to task-relevant signals, effectively mirroring the empirically observed phenomenon that only behaviorally relevant stimuli remain stably retained [98–100].

Moreover, the interpretability afforded by our framework—where each attention weight corresponds to a distinct, memorized spatial patch as well as an immediate visual patch—creates a unique opportunity to perform in silico experiments that mirror microstimulation or lesion studies in NHPs [53, 77]. As demonstrated, directly manipulating self-attention weights in favor of or away from the change location systematically alters detection rates and reaction times. Such controlled perturbations, rarely feasible with standard ViTs, may provide new ways to test causal hypotheses about attention and working memory mechanisms [10, 51, 101].

### 5.5 Broader implications and future directions

This Recurrent ViT opens several avenues for future research. First, richer scenarios—such as multi-object tracking [102,103],dynamic scene understanding [78,104], or tasks requiring mid-trial updates to memorized stimuli [23, 25]—would extend our approach and further test its alignment with primate attentional performance. Incorporating saccadic eye movements, akin to real-world visual search, could allow the model to learn optimal covert and overt strategies in tandem [80, 105]. Additionally, scaling up to deeper, multilayer recurrent architectures may capture the intricate, multi-level feedback loops characteristic of the primate cortex [34,106–108].Second, bridging our Recurrent ViT with reinforcement learning frameworks that incorporate explicit dopamine-like signals [87, 109, 110] could elucidate how value-based attentional modulation emerges in tandem with memory demands. This expansion would dovetail with broader theories of attention, memory, and decision-making as components of a common, computationally grounded process [71,111]. Finally, the interpretability of our approach may inspire future “virtual lesion” or “virtual microstimulation” studies to dissect precisely how feedback, gating, and local competition produce emergent attentional dynamics—a goal shared across computational neuroscience and AI [80,112–116].

### 5.6 Conclusion

Taken together, these findings demonstrate that recapitulating human/NHP-like attention in transformer architectures is possible by introducing explicit top-down influences and recurrent feedback. Our Recurrent ViT significantly narrows the gap between standard, feedforward models of attention [32,33] and the iterative, memory-intensive processes that characterize primate visual cognition [5,15]. By unifying principles of biased competition, working memory, and reward-driven learning within a single framework, we not only advance the biological plausibility of deep vision models but also generate a versatile platform for addressing fundamental questions about how perception, memory, and attention converge to guide adaptive behavior.

## References

[1] Marisa Carrasco. Visual attention: The past 25 years. Vision research, 51(13):1484–1525, 2011.

[2] Kelsey Clark, Ryan Fox Squire, Yaser Merrikhi, and Behrad Noudoost. Visual attention: Linking prefrontal sources to neuronal and behavioral correlates. Progress in neurobiology, 132:59–80, 2015.

[3] James E Hoffman. Visual attention and eye movements. Attention, pages 119–153, 2016.

[4] Roopali Bhatnagar and Jacob L Orquin. A meta-analysis on the effect of visual attention on choice. Journal of Experimental Psychology: General, 151(10):2265, 2022.

[5] Nicole C Rust and Marlene R Cohen. Priority coding in the visual system. Nature Reviews Neuroscience, 23(6):376–388, 2022.

[6] Carrie J McAdams and John HR Maunsell. Effects of attention on the reliability of individual neurons in monkey visual cortex. Neuron, 23(4):765–773, 1999.

[7] Carrie J McAdams and John HR Maunsell. Effects of attention on orientation-tuning functions of single neurons in macaque cortical area v4. Journal of Neuroscience, 19(1):431–441, 1999.

[8] Alexander Thiele and Mark A Bellgrove. Neuromodulation of attention. Neuron, 97(4):769–785, 2018.

[9] Marlene R Cohen and John HR Maunsell. Attention improves performance primarily by reducing interneuronal correlations. Nature neuroscience, 12(12):1594–1600, 2009.

[10] Douglas A Ruff and Marlene R Cohen. Attention increases spike count correlations between visual cortical areas. Journal of Neuroscience, 36(28):7523–7534, 2016.

[11] Michael I Posner, Charles R Snyder, and Brian J Davidson. Attention and the detection of signals. Journal of experimental psychology: General, 109(2):160, 1980.

[12] Edward Awh, Edward K Vogel, and S-H Oh. Interactions between attention and working memory. Neuroscience, 139(1):201–208, 2006.

[13] Adam Gazzaley and Anna C Nobre. Topdown modulation: bridging selective attention and working memory. Trends in cognitive sciences, 16(2):129–135, 2012.

[14] Anastasia Kiyonaga and Tobias Egner. Working memory as internal attention: Toward an integrative account of internal and external selection processes. Psychonomic bulletin & review, 20:228–242, 2013.

[15] Matthew F. Panichello and Timothy J. Buschman. Shared mechanisms underlie the control of working memory and attention. Nature, 592(7855):601–605, 2021.

[16] Klaus Oberauer. Access to information in working memory: exploring the focus of attention. Journal of Experimental Psychology: Learning, Memory, and Cognition, 28(3):411, 2002.

[17] Fiona McNab and Torkel Klingberg. Prefrontal cortex and basal ganglia control access to working memory. Nature neuroscience, 11(1):103– 107, 2008.

[18] Nancy B Carlisle, Jason T Arita, Deborah Pardo, and Geoffrey F Woodman. Attentional templates in visual working memory. Journal of neuroscience, 31(25):9315–9322, 2011.

[19] Dirk van Moorselaar, Jan Theeuwes, and Christian NL Olivers. In competition for the attentional template: Can multiple items within visual working memory guide attention? Journal of Experimental Psychology: Human Perception and Performance, 40(4):1450, 2014.

[20] Nick Berggren and Martin Eimer. Visual working memory load disrupts template-guided attentional selection during visual search. Journal of Cognitive Neuroscience, 30(12):1902–1915, 2018.

[21] Nancy B Carlisle and Árni KristjÁnsson. How visual working memory contents influence priming of visual attention. Psychological Research, 82:833–839, 2018.

[22] Freek van Ede, Sammi R Chekroud, and Anna C Nobre. Human gaze tracks the focusing of attention within the internal space of visual working memory. Journal of Vision, 19(10):133b–133b, 2019.

[23] A Caglar Tas, Steven J Luck, and Andrew Hollingworth. The relationship between visual attention and visual working memory encoding: A dissociation between covert and overt orienting. Journal of experimental psychology: human perception and performance, 42(8):1121, 2016.

[24] Brett Bahle, Valerie M Beck, and Andrew Hollingworth. The architecture of interaction between visual working memory and visual attention. Journal of Experimental Psychology: Human Perception and Performance, 44(7):992, 2018.

[25] Daniela Gresch, Sage EP Boettcher, Freek van Ede, and Anna C Nobre. Shifting attention between perception and working memory. Cognition, 245:105731, 2024.

[26] Mary E Wheeler and Anne M Treisman. Binding in short-term visual memory. Journal of experimental psychology: General, 131(1):48, 2002.

[27] Fabiano Botta and Juan LupiÁñez. Spatial distribution of attentional bias in visuo-spatial working memory following multiple cues. Acta psychologica, 150:1–13, 2014.

[28] Edward Awh, John Jonides, and Patricia A Reuter-Lorenz. Rehearsal in spatial working memory. Journal of Experimental Psychology: Human Perception and Performance, 24(3):780, 1998.

[29] Edward Awh, Lourdes Anllo-Vento, and Steven A Hillyard. The role of spatial selective attention in working memory for locations: Evidence from event-related potentials. Journal of Cognitive Neuroscience, 12(5):840–847, 2000.

[30] Melonie Williams, Pierre Pouget, Leanne Boucher, and Geoffrey F Woodman. Visual– spatial attention aids the maintenance of object representations in visual working memory. Memory & cognition, 41:698–715, 2013.

[31] Sami R Yousif, Monica D Rosenberg, and Frank C Keil. Using space to remember: Shortterm spatial structure spontaneously improves working memory. Cognition, 214:104748, 2021.

[32] Ashish Vaswani, Noam Shazeer, Niki Parmar, Jakob Uszkoreit, Llion Jones, Aidan N Gomez, Lukasz Kaiser, and Illia Polosukhin. Attention is all you need. Advances in neural information processing systems, 30, 2017.

[33] Alexey Dosovitskiy, Lucas Beyer, Alexander Kolesnikov, Dirk Weissenborn, Xiaohua Zhai, Thomas Unterthiner, Mostafa Dehghani, Matthias Minderer, Georg Heigold, Sylvain Gelly, et al. An image is worth 16×16 words: Transformers for image recognition at scale. arXiv preprint 2010.11929, 2020.

[34] Salman Khan, Muzammal Naseer, Munawar Hayat, Syed Waqas Zamir, Fahad Shahbaz Khan, and Mubarak Shah. Transformers in vision: A survey. ACM computing surveys (CSUR), 54(10s):1–41, 2022.

[35] Laurent Itti and Christof Koch. Computational modelling of visual attention. Nature reviews neuroscience, 2(3):194–203, 2001.

[36] Olivier Le Meur, Patrick Le Callet, Dominique Barba, and Dominique Thoreau. A coherent computational approach to model bottom-up visual attention. IEEE transactions on pattern analysis and machine intelligence, 28(5):802– 817, 2006.

[37] Alexander Krü ger, Jan Tü nnermann, and Ingrid Scharlau. Measuring and modeling salience with the theory of visual attention. Attention, Perception, & Psychophysics, 79:1593–1614, 2017.

[38] Jiajie Zou, Yuran Zhang, Jialu Li, Xing Tian, and Nai Ding. Human attention during goal-directed reading comprehension relies on task optimization. Elife, 12:RP87197, 2023.

[39] Paria Mehrani and John K Tsotsos. Selfattention in vision transformers performs perceptual grouping, not attention. arXiv preprint 2303.01542, 2023.

[40] Takuto Yamamoto, Hirosato Akahoshi, and Shigeru Kitazawa. Emergence of human-like attention in self-supervised vision transformers: an eye-tracking study. arXiv preprint 2410.22768, 2024.

[41] Steven J Luck and Edward K Vogel. The capacity of visual working memory for features and conjunctions. Nature, 390(6657):279–281, 1997.

[42] Steven J Luck and Edward K Vogel. Visual working memory capacity: from psychophysics and neurobiology to individual differences. Trends in cognitive sciences, 17(8):391–400, 2013.

[43] Timothy F Brady and Joshua B Tenenbaum. A probabilistic model of visual working memory: Incorporating higher order regularities into working memory capacity estimates. Psychological review, 120(1):85, 2013.

[44] Stephen M Emrich, Holly A Lockhart, and Naseem Al-Aidroos. Attention mediates the flexible allocation of visual working memory resources. Journal of Experimental Psychology: Human Perception and Performance, 43(7):1454, 2017.

[45] Christian NL Olivers, Judith Peters, Roos Houtkamp, and Pieter R Roelfsema. Different states in visual working memory: When it guides attention and when it does not. Trends in cognitive sciences, 15(7):327–334, 2011.

[46] Chunyue Teng and Dwight J Kravitz. Visual working memory directly alters perception. Nature human behaviour, 3(8):827–836, 2019.

[47] Paul M Bays, Sebastian Schneegans, Wei Ji Ma, and Timothy F Brady. Representation and computation in visual working memory. Nature Human Behaviour, pages 1–19, 2024.

[48] Sepp Hochreiter and Jü rgen Schmidhuber. Long short-term memory. Neural computation, 9(8):1735–1780, 1997.

[49] Maximilian Beck, Korbinian Pö ppel Markus Spanring, Andreas Auer, Oleksandra Prudnikova, Michael Kopp, Günter Klambauer, Johannes Brandstetter, and Sepp Hochreiter. xlstm: Extended long short-term memory. arXiv preprint 2405.04517, 2024.

[50] Ramanujan Srinath, Douglas A Ruff, and Marlene R Cohen. Attention improves information flow between neuronal populations without changing the communication subspace. Current Biology, 31(23):5299–5313, 2021.

[51] Tirin Moore and Katherine M Armstrong. Selective gating of visual signals by microstimulation of frontal cortex. Nature, 421(6921):370–373, 2003.

[52] James Cavanaugh and Robert H Wurtz. Subcortical modulation of attention counters change blindness. Journal of Neuroscience, 24(50):11236–11243, 2004.

[53] James Cavanaugh, Bryan D Alvarez, and Robert H Wurtz. Enhanced performance with brain stimulation: attentional shift or visual cue? Journal of Neuroscience, 26(44):11347–11358, 2006.

[54] Robert Egly, Jon Driver, and Robert D Rafal. Shifting visual attention between objects and locations: evidence from normal and parietal lesion subjects. Journal of Experimental Psychology: General, 123(2):161, 1994.

[55] Tormod Thomsen, Karsten Specht, Lars Ersland, and Kenneth Hugdahl. Processing of conflicting cues in an attention-shift paradigm studied with fmri. Neuroscience letters, 380(1-2):138–142, 2005.

[56] Benoit Brisson and Pierre Jolicoeur. Express attentional re-engagement but delayed entry into consciousness following invalid spatial cues in visual search. PLoS One, 3(12):e3967, 2008.

[57] ZL Lu and B Dosher. External noise distinguishes attention mechanisms. Vision research, 38(9):1183 – 1198, 05 1998.

[58] Joshua A. Solomon. The effect of spatial cues on visual sensitivity. Vision Research, 44(12):1209– 1216, 2004.

[59] E.Leslie Cameron, Joanna C Tai, and Marisa Carrasco. Covert attention affects the psychometric function of contrast sensitivity. Vision Research, 42(8):949–967, 2002.

[60] Hermann J Mü ller and John M Findlay. Sensitivity and criterion effects in the spatial cuing of visual attention. Perception & Psychophysics, 42(4):383–399, 1987.

[61] Harold L Hawkins, Steven A Hillyard, Steven J Luck, Mustapha Mouloua, Cathryn J Downing, and Donald P Woodward. Visual attention modulates signal detectability. Journal of Experimental Psychology: Human Perception and Performance, 16(4):802, 1990.

[62] IJ Saltzman and WR Garner. Reaction time as a measure of span of attention. The Journal of psychology, 25(2):227–241, 1948.

[63] Jerry S Carlson, C Mark Jensen, and Keith F Widaman. Reaction time, intelligence, and attention. Intelligence, 7(4):329–344, 1983.

[64] William Prinzmetal, Christin McCool, and Samuel Park. Attention: reaction time and accuracy reveal different mechanisms. Journal of Experimental Psychology: General, 134(1):73, 2005.

[65] Deborah A Jehu, Alyssa Desponts, Nicole Paquet, and Yves Lajoie. Prioritizing attention on a reaction time task improves postural control and reaction time. International Journal of Neuroscience, 125(2):100–106, 2015.

[66] James P. Herman and Richard J. Krauzlis. Color-Change Detection Activity in the Primate Superior Colliculus. eNeuro, 4(2):ENEURO.0046– 17.2017, 3 2017.

[67] Geoffrey M. Ghose and John H. R. Maunsell. Attentional modulation in visual cortex depends on task timing. Nature, 419(6907):616–620, 2002.

[68] Jitendra Sharma, Hiroki Sugihara, Yarden Katz, James Schummers, Joshua Tenenbaum, and Mriganka Sur. Spatial Attention and Temporal Expectation Under Timed Uncertainty Predictably Modulate Neuronal Responses in Monkey V1. Cerebral Cortex, 25(9):2894–2906, 2015.

[69] Santiago Jaramillo and Anthony M Zador. The auditory cortex mediates the perceptual effects of acoustic temporal expectation. Nature Neuroscience, 14(2):246–251, 2011.

[70] Anna C. Nobre and Freek van Ede. Anticipated moments: temporal structure in attention. Nature Reviews Neuroscience, 19(1):34–48, 2018.

[71] Eric I Knudsen. Fundamental components of attention. Annu. Rev. Neurosci., 30(1):57–78, 2007.

[72] Farhan Baluch and Laurent Itti. Mechanisms of top-down attention. Trends in neurosciences, 34(4):210–224, 2011.

[73] Jillian H Fecteau and Douglas P Munoz. Salience, relevance, and firing: a priority map for target selection. Trends in cognitive sciences, 10(8):382–390, 2006.

[74] Jacqueline Gottlieb and Puiu Balan. Attention as a decision in information space. Trends in Cognitive Sciences, 14(6):240–248, 2010.

[75] James W Bisley and Michael E Goldberg. Attention, Intention, and Priority in the Parietal Lobe. Annual review of neuroscience, 33(1):1 – 21, 2010.

[76] Jeremy M Wolfe. Guided search 6.0: An updated model of visual search. Psychonomic Bulletin & Review, 28(4):1060–1092, 2021.

[77] Koorosh Mirpour, Wei Song Ong, and James W Bisley. Microstimulation of posterior parietal cortex biases the selection of eye movement goals during search. Journal of neurophysiology, 104(6):3021–3028, 2010.

[78] Dominique Lamy, Hannah Segal, and Lital Ruderman. Grouping does not require attention. Perception & Psychophysics, 68(1):17–31, 2006.

[79] Robert Desimone, John Duncan, et al. Neural mechanisms of selective visual attention. Annual review of neuroscience, 18(1):193–222, 1995.

[80] Richard J Krauzlis, Lee P Lovejoy, and Alexandre Zénon. Superior colliculus and visual spatial attention. Annual review of neuroscience, 36:165– 182, 2013.

[81] Michael A Silver, David Ress, and David J Heeger. Topographic maps of visual spatial attention in human parietal cortex. Journal of neurophysiology, 94(2):1358–1371, 2005.

[82] Rafiq Huda, Grayson O Sipe, Vincent Breton-Provencher, K Guadalupe Cruz, Gerald N Pho, Elie Adam, Liadan M Gunter, Austin Sullins, Ian R Wickersham, and Mriganka Sur. Distinct prefrontal top-down circuits differentially modulate sensorimotor behavior. Nature communications, 11(1):6007, 2020.

[83] Andrew D Bolton, Yasunobu Murata, Rory Kirchner, Sung-Yon Kim, Andrew Young, Tru Dang, Yuchio Yanagawa, and Martha Constantine-Paton. A diencephalic dopamine source provides input to the superior colliculus, where d1 and d2 receptors segregate to distinct functional zones. Cell reports, 13(5):1003–1015, 2015.

[84] Kamil Pradel, Gniewosz Drwiga, and Tomasz Błasiak. Superior colliculus controls the activity of the rostromedial tegmental nuclei in an asymmetrical manner. Journal of Neuroscience, 41(18):4006–4022, 2021.

[85] Jaclyn Essig and Gidon Felsen. Warning! dopaminergic modulation of the superior colliculus. Trends in neurosciences, 39(1):2–4, 2016.

[86] Juan Pérez-FernÁndez, Andreas A Kardamakis, Daichi G Suzuki, Brita Robertson, and Sten Grillner. Direct dopaminergic projections from the snc modulate visuomotor transformation in the lamprey tectum. Neuron, 96(4):910–924, 2017.

[87] Okihide Hikosaka, Kae Nakamura, and Hiroyuki Nakahara. Basal ganglia orient eyes to reward. Journal of neurophysiology, 95(2):567– 584, 2006.

[88] Clayton Hickey, Leonardo Chelazzi, and Jan Theeuwes. Reward changes salience in human vision via the anterior cingulate. Journal of Neuroscience, 30(33):11096–11103, 2010.

[89] Michel Failing and Jan Theeuwes. Selection history: How reward modulates selectivity of visual attention. Psychonomic bulletin & review, 25(2):514–538, 2018.

[90] Karl Friston, James Kilner, and Lee Harrison. A free energy principle for the brain. Journal of physiology-Paris, 100(1-3):70–87, 2006.

[91] Harriet Feldman and Karl J Friston. Attention, uncertainty, and free-energy. Frontiers in human neuroscience, 4:215, 2010.

[92] Karl J Friston, Tamara Shiner, Thomas FitzGerald, Joseph M Galea, Rick Adams, Harriet Brown, Raymond J Dolan, Rosalyn Moran, Klaas Enno Stephan, and Sven Bestmann. Dopamine, affordance and active inference. PLoS computational biology, 8(1):e1002327, 2012.

[93] Bijan Khezri. Free energy principle (fep). In Governing Continuous Transformation: Re-framing the Strategy-Governance Conversation, pages 33–41. Springer, 2022.

[94] Pietro Mazzaglia, Tim Verbelen, Ozan Çatal, and Bart Dhoedt. The free energy principle for perception and action: A deep learning perspective. Entropy, 24(2):301, 2022.

[95] Mohammed Hassanin, Saeed Anwar, Ibrahim Radwan, Fahad Shahbaz Khan, and Ajmal Mian. Visual attention methods in deep learning: An indepth survey. Information Fusion, 108:102417, 2024.

[96] John H Reynolds, Leonardo Chelazzi, and Robert Desimone. Competitive mechanisms subserve attention in macaque areas v2 and v4. Journal of Neuroscience, 19(5):1736–1753, 1999.

[97] James W Bisley and Koorosh Mirpour. The neural instantiation of a priority map. Current opinion in psychology, 29:108–112, 2019.

[98] Robert Desimone. Neural mechanisms for visual memory and their role in attention. Proceedings of the National Academy of Sciences, 93(24):13494–13499, 1996.

[99] Stefan Van der Stigchel, Artem V Belopolsky, Judith C Peters, Jasper G Wijnen, Martijn Meeter, and Jan Theeuwes. The limits of top-down control of visual attention. Acta psychologica, 132(3):201–212, 2009.

[100] Tae-Ho Lee, Steven G Greening, and Mara Mather. Encoding of goal-relevant stimuli is strengthened by emotional arousal in memory. Frontiers in psychology, 6:1173, 2015.

[101] James P Herman, Fabrice Arcizet, and Richard J Krauzlis. Attention-related modulation of caudate neurons depends on superior colliculus activity. eLife, 9:e53998, sep 2020.

[102] Katherine C Bettencourt and David C Somers. Effects of target enhancement and distractor suppression on multiple object tracking capacity. Journal of vision, 9(7):9–9, 2009.

[103] Hauke S Meyerhoff, Frank Papenmeier, and Markus Huff. Studying visual attention using the multiple object tracking paradigm: A tutorial review. Attention, Perception, & Psychophysics, 79:1255–1274, 2017.

[104] Jeremy M Wolfe, Melissa L-H Võ, Karla K Evans, and Michelle R Greene. Visual search in scenes involves selective and nonselective pathways. Trends in cognitive sciences, 15(2):77–84, 2011.

[105] Priyanka Gupta and Devarajan Sridharan. Presaccadic attention does not facilitate the detection of changes in the visual field. PLoS Biology, 22(1):e3002485, 2024.

[106] Daniel J Felleman and David C Van Essen. Distributed hierarchical processing in the primate cerebral cortex. Cerebral cortex (New York, NY: 1991), 1(1):1–47, 1991.

[107] Tim C Kietzmann, Courtney J Spoerer, Lynn KA Sö rensen Radoslaw M Cichy, Olaf Hauk, and Nikolaus Kriegeskorte. Recurrence is required to capture the representational dynamics of the human visual system. Proceedings of the National Academy of Sciences, 116(43):21854–21863, 2019.

[108] Chengxu Zhuang, Siming Yan, Aran Nayebi, Martin Schrimpf, Michael C Frank, James J DiCarlo, and Daniel LK Yamins. Unsupervised neural network models of the ventral visual stream. Proceedings of the National Academy of Sciences, 118(3):e2014196118, 2021.

[109] Matthew Botvinick, Jane X Wang, Will Dabney, Kevin J Miller, and Zeb Kurth-Nelson. Deep reinforcement learning and its neuroscientific implications. Neuron, 107(4):603–616, 2020.

[110] Benedicte M Babayan, Naoshige Uchida, and Samuel J Gershman. Belief state representation in the dopamine system. Nature communications, 9(1):1891, 2018.

[111] Ilya E Monosov. How outcome uncertainty mediates attention, learning, and decision-making. Trends in neurosciences, 43(10):795–809, 2020.

[112] Valerio Mante, David Sussillo, Krishna V Shenoy, and William T Newsome. Context-dependent computation by recurrent dynamics in prefrontal cortex. nature, 503(7474):78–84, 2013.

[113] Ricardo Gattass and Robert Desimone. Effect of microstimulation of the superior colliculus on visual space attention. Journal of cognitive neuroscience, 26(6):1208–1219, 2014.

[114] Thomas Miconi and Rufin VanRullen. A feedback model of attention explains the diverse effects of attention on neural firing rates and receptive field structure. PLoS computational biology, 12(2):e1004770, 2016.

[115] Guoyang Liu, Jindi Zhang, Antoni B Chan, and Janet H Hsiao. Human attention guided explainable artificial intelligence for computer vision models. Neural Networks, 177:106392, 2024.

[116] Giuseppe Cartella, Marcella Cornia, Vittorio Cuculo, Alessandro D’Amelio, Dario Zanca, Giuseppe Boccignone, and Rita Cucchiara. Trends, applications, and challenges in human attention modelling. arXiv preprint 2402.18673, 2024.

