## supplement for "A recurrent vision transformer shows signatures of primate visual attention"

immediate

February 13, 2025

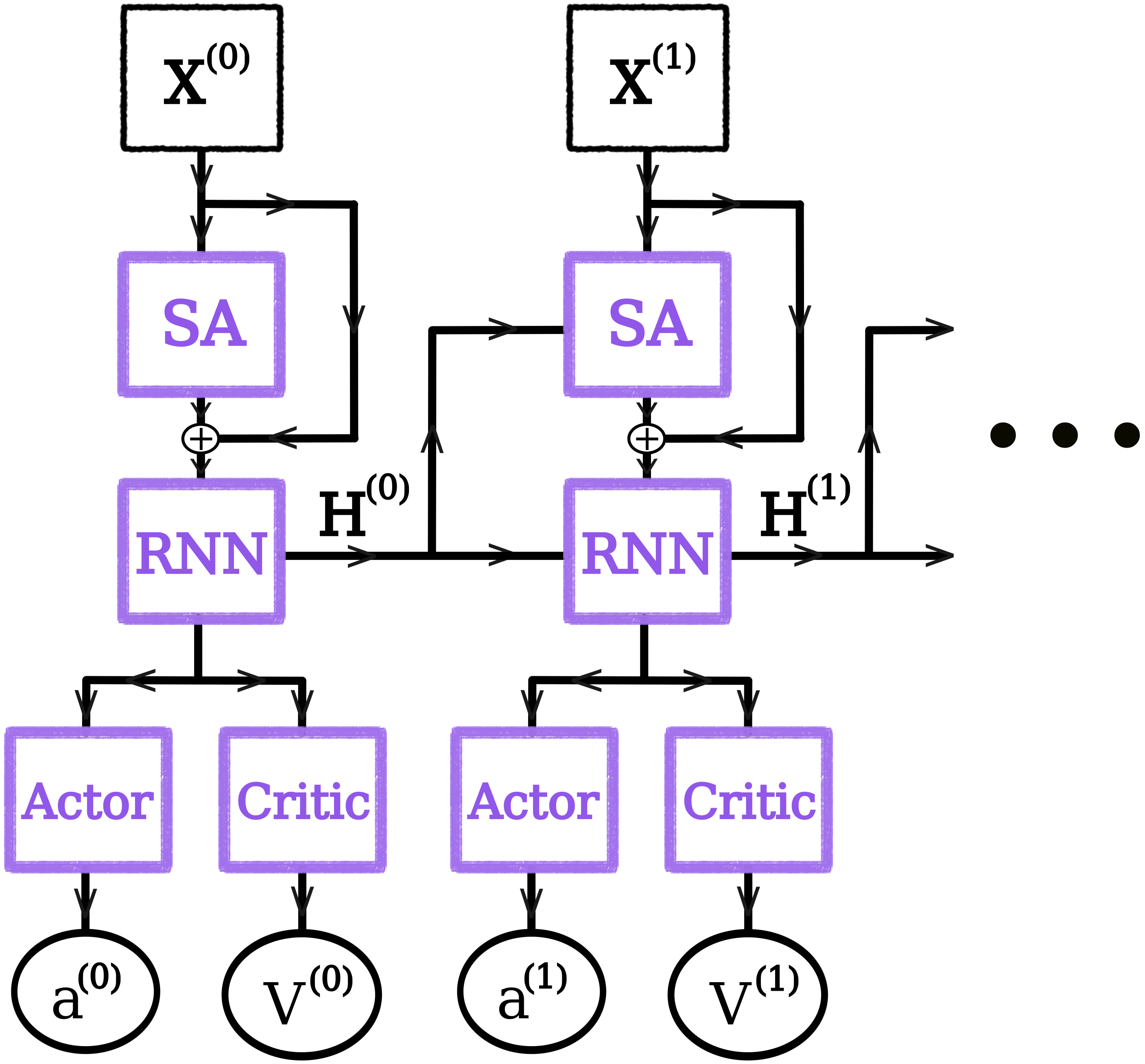

**Figure 1.** Model Concept. Arrows indicate directional flow of information and processing. Images $(X^{(t)})$ are generated from the environment. These images are then fed through the Recurrent ViT from which the resulting output is passed to an RNN. The internal state of the RNN is updated used in the decision making process, and also passed to the Recurrent ViT and RNN at the next time step. Purple boxes denote artificial neural networks

### 1 Methods

#### 1.1 Model Overview

The objective of the model is to utilize immediate visual inputs in order to update an internal state with sufficient immediate and past visual information such that downstream decoders can estimate value and take action. We utilize a self-attention (SA) mechanism to construct the visual percept used to update the internal state of the RNN. Self-attention is computed based on the immediate visual inputs and feedback from the RNN. The following sections will describe the motivations and details of this process in more depth.

#### 1.2 Pre-Processing, Content Selection, and Construction for Visual Working Memory

Given a visual scene (an image) of dimension $H \times W \times C$, denoted by $\mathcal{O}^{(t)} \in \mathbb{R}^{H \times W \times C}$ at time $t$, the agent views the entire scene through a fixation at center field. During preprocessing, the image is partitioned into a set of patches, $\{\mathbf{o_i}^{(t)}\}_{i=1}^{n_{patch}}$, where each patch is of size $H_{patch} \times W_{patch} \times C_{patch}$. The original visual patches are then transformed into a compact set of internal representations, $\{\mathbf{x_i}^{(t)}\}_{i=1}^{n_{patch}}$, each $\mathbf{x_i}^{(t)} \in \mathbb{R}^{H_{patch} \times W_{patch} \times C_{patch}}$ and typically satisfying

$$\dim(\mathbf{x_i}^{(t)}) \ll \dim\big(\mathbf{o_i}^{(t)}\big).$$

Together, the collection of these feature patches forms the immediate visual information available to the agent at time $t$, denoted by $\mathbf{X}^{(t)} = \{\mathbf{x_i}^{(t)}\}_{i=1}^{n_{patch}}$.

##### 1.3 Spatially Oriented Visual Working Memory

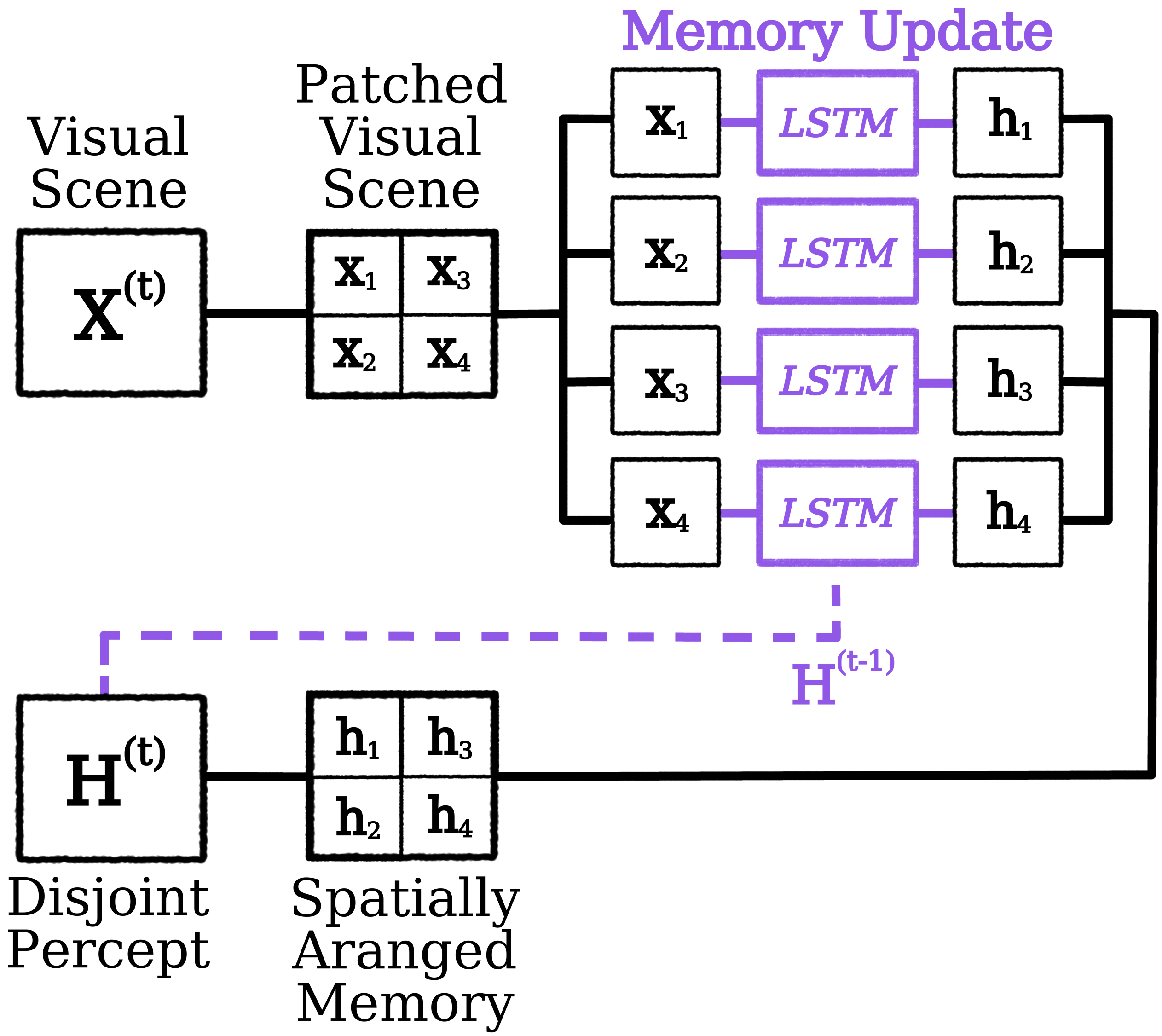

**Figure 2.** The patch-based LSTM. In essence, it is a standard LSTM architecture [3, 11]. The only difference is that we apply the LSTM in parallel to all patches in a visual scene. This yields separate recurrent states for each patch. However, the weights local to the LSTM are shared among all patches. I.e., the weights utilized to construct $h_4^{(t)}$ given $x_4^{(t)}$ and $h_4^{(t-1)}$ are the same weights sued to construct $h_3^{(t)}$ given $x_3^{(t)}$ and $h_3^{(t-1)}$ and the two operations occur in parallel. The patch-based LSTM does not mix information among input patches.

Following numerous experimental findings, we allow our model to maintain a spatially arranged visual working memory [22, 23, 27, 29], in which each patch location $i$ has a corresponding patched memory component $\mathbf{c_i}^{(t)} \in \mathbb{R}^{d_{mem}}$ within the RNN. For updating the internal state of our RNN, we utilize the operations and functions described in the LSTM architecture [3, 11]. For the remainder of our description, we will refer to the collection $C^{(t)} = \{c_i^{(t)}\}_{i=1}^{n_{patch}}$ the VWM state and $c_i^{(t)}$ a VWM patch, where

$$\mathbf{c_i}^{(t)} = \mathbf{f_i}^{(t)} \odot \mathbf{c_i}^{(t-1)} + \mathbf{u_i}^{(t)} \odot \boldsymbol{\psi_i}^{(t)}, \tag{1}$$

where

$$\mathbf{f_i}^{(t)} = F\big(\mathbf{x_i}^{(t)}, \mathbf{h_i}^{(t-1)}\big), \quad \mathbf{u_i}^{(t)} = U\big(\mathbf{x_i}^{(t)}, \mathbf{h_i}^{(t-1)}\big), \quad \boldsymbol{\psi_i}^{(t)} = \Psi\big(\mathbf{x_i}^{(t)}, \mathbf{h_i}^{(t-1)}\big).$$

The operator $\odot$ denotes elementwise multiplication. Here, $\mathbf{f_i}^{(t)} \in [0,1]^{d_{mem}}$ determines which parts of $\mathbf{c_i}^{(t-1)}$ are *forgotten* (i.e., decayed), while $\mathbf{u_i}^{(t)} \in [-1,1]^{d_{mem}}$ and $\boldsymbol{\psi_i}^{(t)} \in \mathbb{R}^{d_{mem}}$ selectively modulate and propose new content. Altogether, these operations enable dynamic insertion, maintenance, and forgetting of information in $\mathbf{c_i}^{(t)}$.

The activated memory patch, $h_i^{(t)}$ is constructed from the VWM patch, $c_i^{(t)}$:

$$\mathbf{h_i}^{(t)} = \boldsymbol{\phi_i}^{(t)} \odot \left(\frac{\mathbf{c_i}^{(t)}}{\mathbf{n_i}^{(t)}}\right),$$

where $\boldsymbol{\phi_i}^{(t)} = \Phi\big(\mathbf{x_i}^{(t)}, \mathbf{h_i}^{(t-1)}\big) \in [0,1]^{d_{mem}}$ selects elements of $\mathbf{c_i}^{(t)}$ for downstream processing, and $\mathbf{n_i}^{(t)}$ is a normalization term. Each function $F, U, \Psi, \Phi$ is parameterized by a feedforward neural network that receives $\mathbf{x_i}^{(t)}$ and $\mathbf{h_i}^{(t-1)}$ as inputs. Although the parameters of these functions remain fixed once trained, the recurrent operation through $\mathbf{h_i}^{(t)}$ allows the system to track temporal dynamics.

#### 1.4 A Disjoint Memory

Following the ideas presented by Knudsen [13], the activated subset of working memeory is central for decision-making and planning. However, a patched RNN as described above presents a clear shortcoming: each $\mathbf{h_i}^{(t)}$ encodes the content of its own patch independently, without explicit awareness of neighboring patches. If downstream processes (*e.g.*, a decoder $\pi$) require spatial or contextual relationships among patches, they must construct these relationships entirely from the population of VWM patches. Moreover, the problem intensifies over time. If the network must integrate information from patches across multiple timesteps (e.g., $\mathbf{x_i}^{(\tau)}$ and $\mathbf{x_j}^{(\tau)}$ for $\tau < t$), then each VWM patch must *retain* all potentially significant current and past features useful for task-relevant decoding by downstream networks. This approach quickly becomes intractable, as it demands that the architecture store a large number of unique *conjunctions* of spatio-temporal features. This challenge aligns with the combinatorial explosion recognized by Tsotsos [26] as a core difficulty in perceptual organization.

To circumvent these limitations, we introduce self-attention into the encoding process, encouraging each patch's representation to reflect the context provided by the other patches within the same timestep. In doing so, we create a spatially integrated or *context-aware* activated memory before the information even updates the VWM.

##### 1.5 Self-attention and spatially aware activated memory

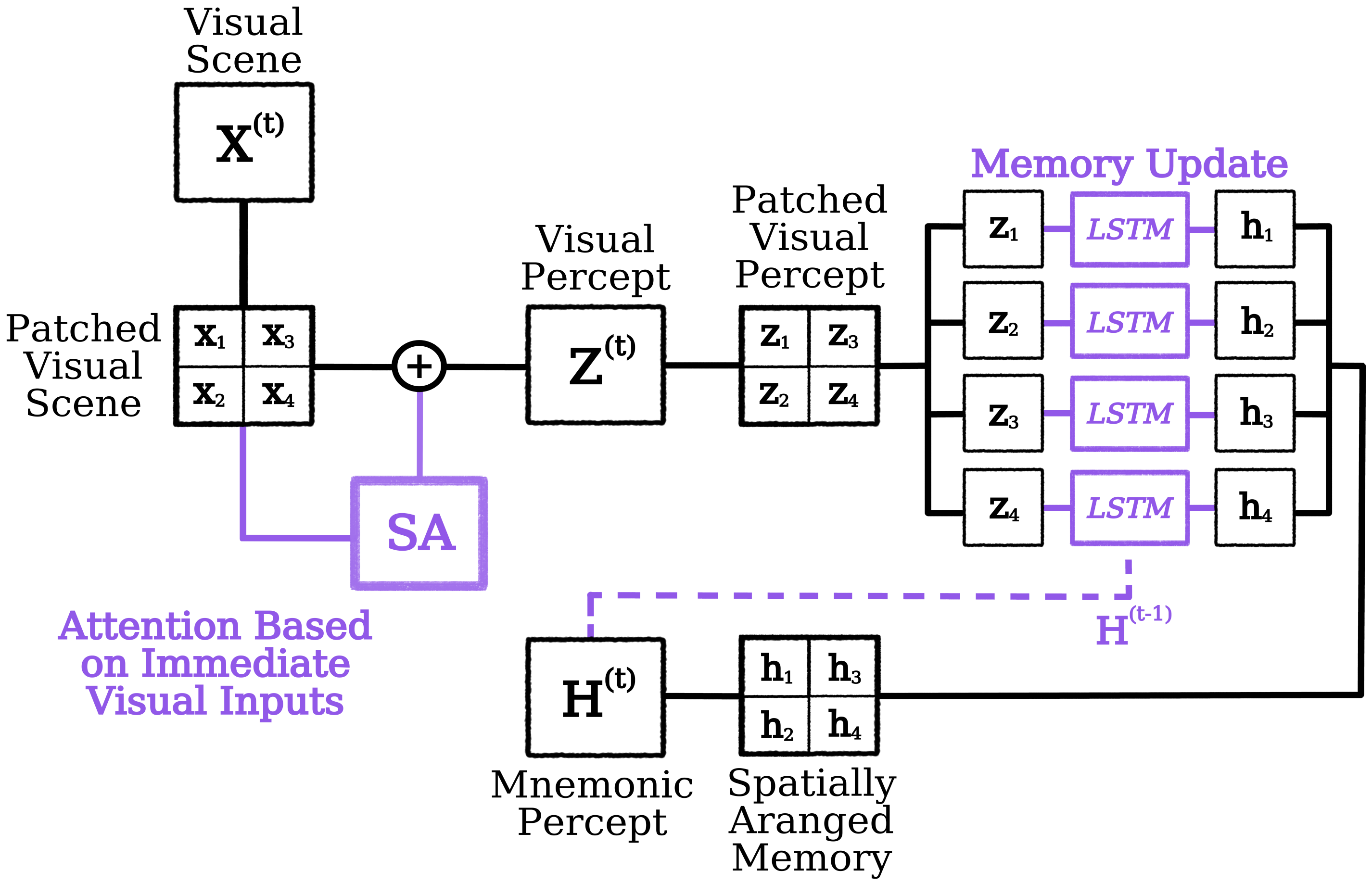

**Figure 3.** The self-attention mechanism merges immediate visual inputs with spatial context. This allows the patch-based LSTM to store relevant visual information form patches and their potential significance given the content in the visual scene.

We wish to obtain a scene-level representation $\mathbf{Z}^{(t)} = \{\mathbf{z_i}^{(t)}\}_{i=1}^{n_{patch}}$ such that each $\mathbf{z_i}^{(t)}$ encodes the task-relevant spatial relationships among the visual feature patches $\mathbf{x_1}^{(t)}, \ldots, \mathbf{x_{n_{patch}}}^{(t)}$. By doing so, the patch-based LSTM will be able to utilize immediate visual information within a patch and task-relevant spatial context to update internal states. Formally, we want

$$\mathbf{z_i}^{(t)} = f_z\Big(\mathbf{x_i}^{(t)}, \{\mathbf{x_j}^{(t)}\}_{j \neq i}\Big),$$

A straightforward way to implement this is via self-attention:

$$\mathbf{z_i}^{(t)} = \mathbf{x_i}^{(t)} + \sum_{j=1}^{n_{patch}} a_{ij}^{(t)} \, \mathbf{v_j}^{(t)}, \tag{2}$$

where $\mathbf{v_j}^{(t)} = V\big(\mathbf{x_j}^{(t)}\big)$ is a function that maps the feature patch into a latent space and $a_{ij}^{(t)} = A\big(\mathbf{x_i}^{(t)}, \mathbf{x_j}^{(t)}\big)$ indicates the *relative importance* of $\mathbf{x_j}^{(t)}$ with respect to $\mathbf{x_i}^{(t)}$. To ensure a proper probability-like weighting, we impose $\sum_{j=1}^{n_{patch}} a_{ij}^{(t)} = 1$ with $a_{ij}^{(t)} \in (0, 1)$. A typical choice for $a_{ij}^{(t)}$ is:

$$a_{ij}^{(t)} \;=\; \frac{\exp\Big(\big\langle \mathbf{q_i}^{(t)}, \mathbf{k_j}^{(t)} \big\rangle\Big)}{\sum_{m=1}^{n_{patch}} \exp\Big(\big\langle \mathbf{q_i}^{(t)}, \mathbf{k_m}^{(t)} \big\rangle\Big)},$$

where $\mathbf{q_i}^{(t)} = Q(\mathbf{x_i}^{(t)})$ and $\mathbf{k_j}^{(t)} = K(\mathbf{x_j}^{(t)})$ are *query* and *key* functions, respectively. Interpreting $a_{ij}^{(t)}$ as a salient feature map has strong parallels to the saliency map hypothesis [14]; however, we adopt a *winner-takes-most* approach rather than a strict winner-takes-all (WTA), common in many self-attention applications. In principle, should $a_{i,j^*}^{(t)} \approx 1$ for some $j^*$ and $a_{i,m}^{(t)} \approx 0$ for $m \neq j^*$, we recover WTA-like mechanism.

After computing $\mathbf{Z}^{(t)}$ for the entire scene, the visual percept patches, $z_i^{(t)}$, are used to update the VWM patches:

$$\mathbf{c_i}^{(t)} = \mathbf{f_i}^{(t)} \odot \mathbf{c_i}^{(t-1)} + \mathbf{u_i}^{(t)} \odot \boldsymbol{\psi_i}^{(t)},$$

where each function is now evaluated using $\mathbf{z_i}^{(t)}$ rather than the immediate visual scene patch in isolation, $\mathbf{x_i}^{(t)}$. This approach solves the *spatial* integration problem in the current timestep. Yet, any feature with contextual importance *across* timesteps remains challenging: we still need a mechanism to capture top-down feedback or *memory-based* salience.

#### 1.6 Recurrent Feedback From Memory

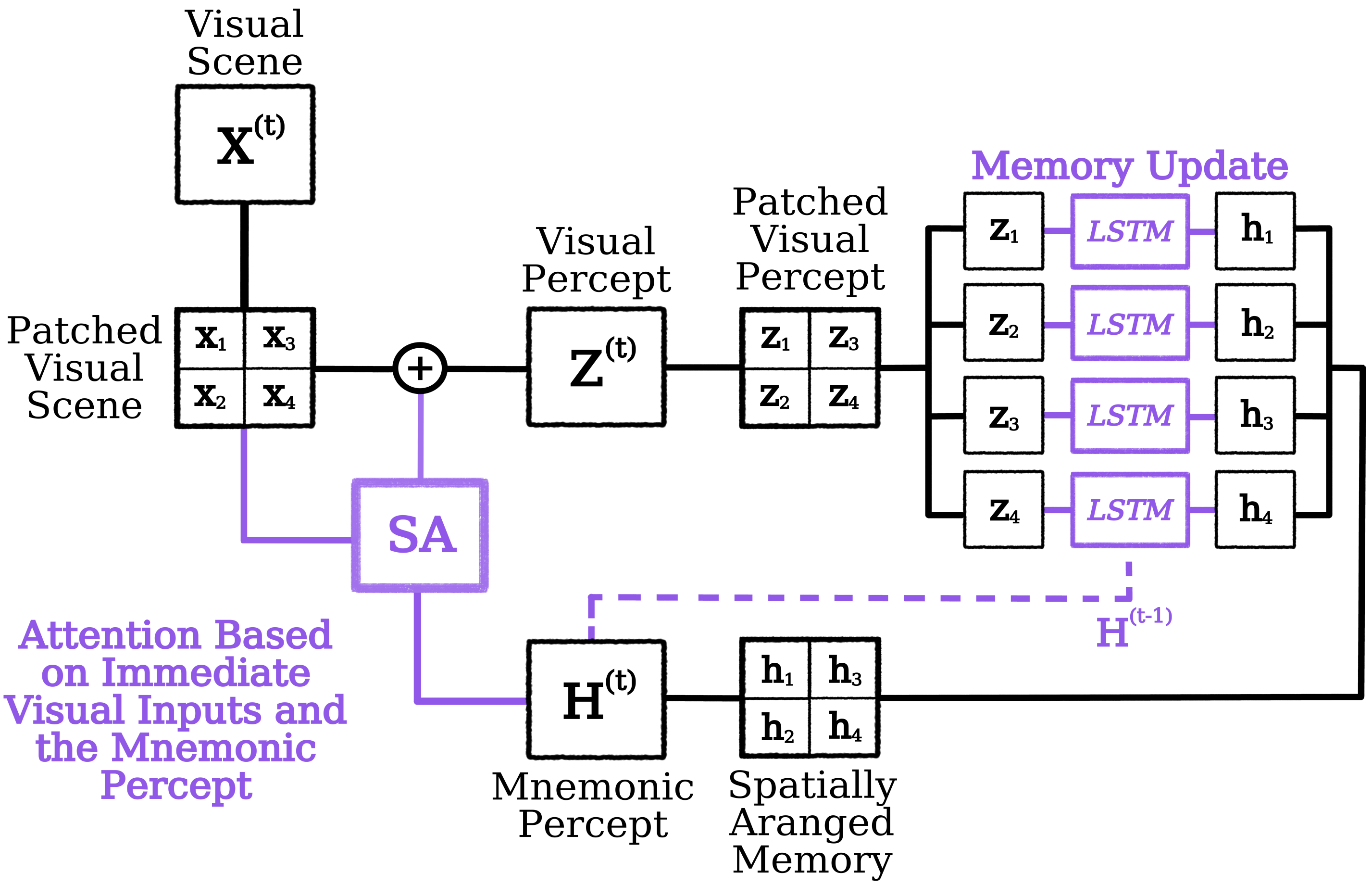

**Figure 4.** A depiction of recurrent self-attention. In order to construct a meaningful representation of the current visual scene, spatial and temporal context are merged with immediate visual inputs.

Knudsen [13] describes a feedback loop in which working memory provides top-down signals that bias neural representations relevant to the organism's current goals. In the context of the biased competition model [6], working memory holds an *attentional template* that biases competition in favor of task-relevant representations. However, in practice it is not clear how this mnemonic feedback is/should be implemented. In this we simplify (and constrain) the problem to implementing recurrent feedback from the patch-based LSTM to the self-attention mechanism of the ViT. Hence, we evaluate three different methods in terms of their ability to yield primate-like behavior signatures of attention. We call the vision transformer with mnemonic feedback the recurrent ViT.

#### 1.7 Mnemonic Guidance

##### 1.7.1 Visual working memory as tokens

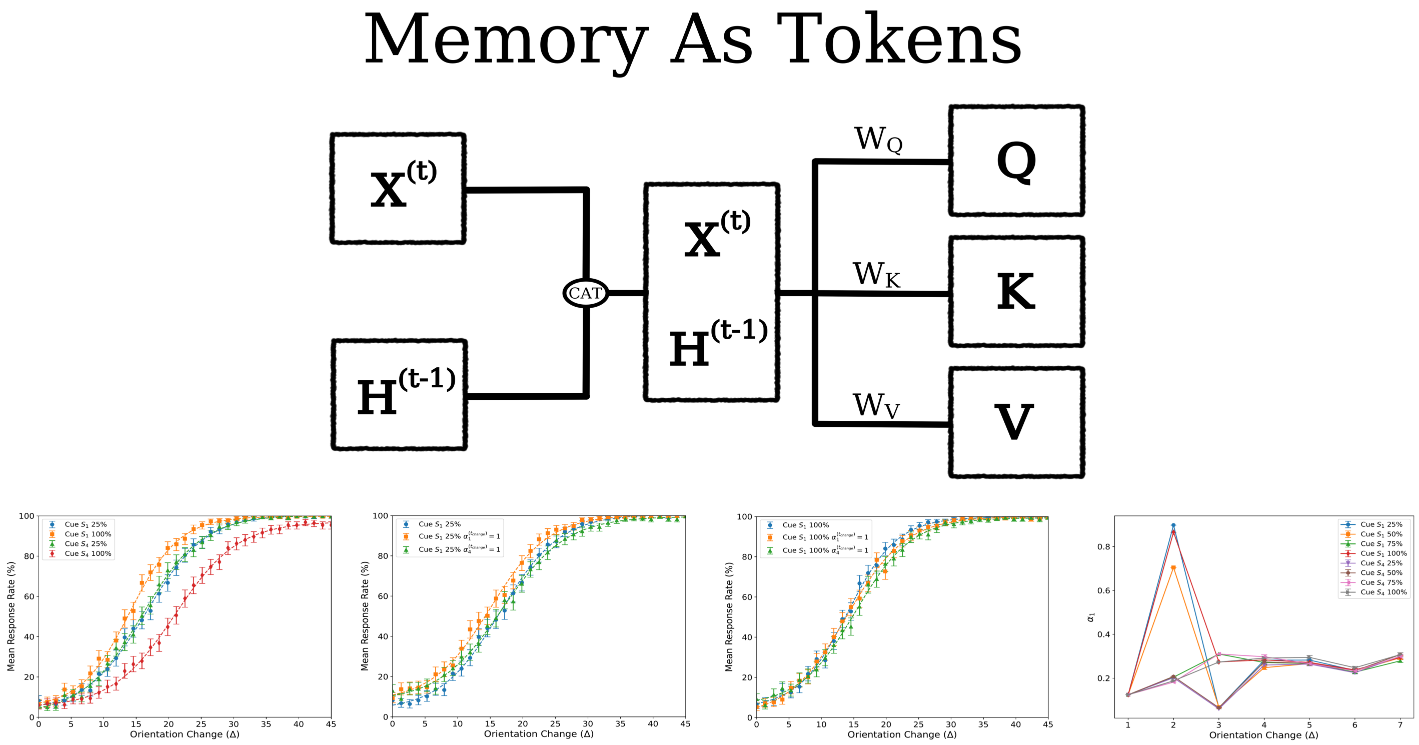

**Figure 5.** Circuit diagram and some behavioral results for a model in which recurrent feedback is implemented via concatenating recurrent states to immediate visual inputs. The concatenation effectively doubles the number of patches in the visual percept $Z^{(t)}$, but we reduce the number back to $n_{patch}$ before the transmission of the visual percept to the LSTM. While we see cue effects on behavior, this model lacks any significant attention modulation effects. In addition, we also observe that the cue has little effect on downstream deployment of attention at later time points.

The first recurrent feedback method we evaluate is one in which we concatenate the mnemonic percept to the visual input. Thus, the input to the self-attention mechanism is

$$\tilde{\mathbf{X}} = Concatenate[\mathbf{X}^{(t)}, \mathbf{H}^{(t-1)}]$$

where $\tilde{\mathbf{X}} \in \mathbb{R}^{2n_{patch}, d_{model}}$. From here we define:

$$\mathbf{q}^{(t)}_{\tilde{X},i} = Q_{\tilde{X}}\big(\tilde{\mathbf{x}}^{(t)}_i\big), \quad \mathbf{k}^{(t)}_{\tilde{X},j} = K_{\tilde{X}}\big(\tilde{\mathbf{x}}^{(t)}_j\big), \quad \mathbf{v}^{(t)}_{\tilde{X},j} = V_{\tilde{X}}\big(\tilde{\mathbf{x}}^{(t)}_j\big),$$

The attention weights are given by

$$\alpha^{(t)}_{i,j} = \frac{\exp\Big(\langle \mathbf{q}^{(t)}_{\tilde{X},i},\ \mathbf{k}^{(t)}_{\tilde{X},j}\rangle\Big)}{\sum_{m=1}^{n_{\text{patch}}} \exp\Big(\langle \mathbf{q}^{(t)}_{\tilde{X},i},\ \mathbf{k}^{(t)}_{\tilde{X},m}\rangle\Big)}. \tag{3}$$

We then compute the output representation as

$$\mathbf{z}_i^{(t)} = \mathbf{x}_i^{(t)} + \sum_{j=1}^{2n_{\text{patch}}} \alpha_{i,j}^{(t)} \mathbf{v}_{\tilde{X},j}^{(t)}. \tag{4}$$

Here, we only take the first $n_{patch}$ entries $\{\mathbf{z}_i^{(t)}\}_{i=1}^{n_{patch}}$. The reason for this is because there are only $n_{patch}$ recurrent states in the patch-based LSTM, and $Z^{(t)} = \{\mathbf{z}^{(t)}\}_{i=1}^{n_{patch}}$ is only used as the input to the LSTM.

##### 1.7.2 Additive Feedback from Visual Working Memory

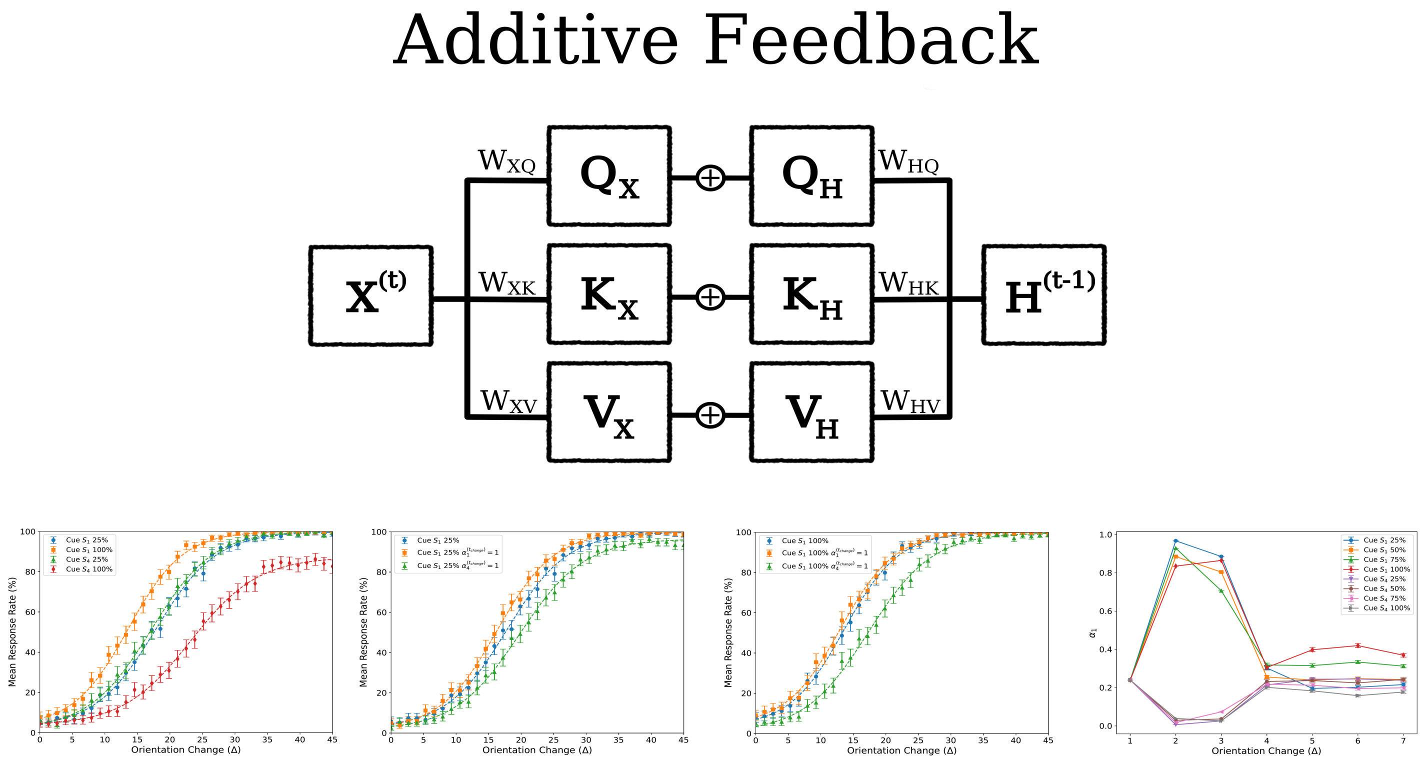

**Figure 6.** Circuit diagram and some behavioral results for a model in which recurrent feedback is implemented via the addition of parallel projections onto the Q, K, V self-attention components. There are significant cue effects and slight attention modulation effects on behavior. The cue also affects the deployment of attention at the time of change, albeit only slightly.

We split the standard self-attention operation into two parallel pathways: one for the bottom-up immediate visual inputs, $\mathbf{x}_i^{(t)}$, and one for the top-down mnemonic inputs, $\mathbf{h}_i^{(t)}$. Define:

$$\mathbf{q}_{X,i}^{(t)} = Q_X\big(\mathbf{x}_i^{(t)}\big), \quad \mathbf{k}_{X,j}^{(t)} = K_X\big(\mathbf{x}_j^{(t)}\big), \quad \mathbf{v}_{X,j}^{(t)} = V_X\big(\mathbf{x}_j^{(t)}\big),$$

and

$$\mathbf{q}_{H,i}^{(t)} = Q_H\big(\mathbf{h}_i^{(t)}\big), \quad \mathbf{k}_{H,j}^{(t)} = K_H\big(\mathbf{h}_j^{(t)}\big), \quad \mathbf{v}_{H,j}^{(t)} = V_H\big(\mathbf{h}_j^{(t)}\big).$$

The attention weights are given by

$$\alpha_{i,j}^{(t)} = \frac{\exp\Big(\langle \mathbf{q}_{X,i}^{(t)} + \mathbf{q}_{H,i}^{(t)},\ \mathbf{k}_{X,j}^{(t)} + \mathbf{k}_{H,j}^{(t)} \rangle\Big)}{\sum_{m=1}^{n_{\text{patch}}} \exp\Big(\langle \mathbf{q}_{X,i}^{(t)} + \mathbf{q}_{H,i}^{(t)},\ \mathbf{k}_{X,m}^{(t)} + \mathbf{k}_{H,m}^{(t)} \rangle\Big)}. \tag{5}$$

We then compute the output representation as

$$\mathbf{z}_i^{(t)} \;=\; \mathbf{x}_i^{(t)} \;+\; \sum_{j=1}^{n_{\text{patch}}} \alpha_{i,j}^{(t)}\Big(\mathbf{v}_{X,j}^{(t)} \;+\; \mathbf{v}_{H,j}^{(t)}\Big). \tag{6}$$

In this additive design, features from the visual inputs and the mnemonic percept patches $\{\mathbf{h}_i^{(t)}\}$ both contribute to the self-attention mechanism by modifying the inner product in the numerator of (5) and by merging the corresponding values in (6).

##### 1.7.3 Multiplicative Feedback from Visual Working Memory

To incorporate multiplicative feedback, we instead define:

$$\alpha_{i,j}^{(t)} \;=\; \frac{\exp\Big(\langle \mathbf{q}_{X,i}^{(t)} \odot \mathbf{q}_{H,i}^{(t)},\; \mathbf{k}_{X,j}^{(t)} \odot \mathbf{k}_{H,j}^{(t)} \rangle\Big)}{\sum_{m=1}^{n_{\text{patch}}} \exp\Big(\langle \mathbf{q}_{X,i}^{(t)} \odot \mathbf{q}_{H,i}^{(t)},\; \mathbf{k}_{X,m}^{(t)} \odot \mathbf{k}_{H,m}^{(t)} \rangle\Big)}, \tag{7}$$

and

$$\mathbf{z}_i^{(t)} \;=\; \mathbf{x}_i^{(t)} \;+\; \sum_{j=1}^{n_{\text{patch}}} \alpha_{i,j}^{(t)}\Big(\mathbf{v}_{X,j}^{(t)} \;\odot\; \mathbf{v}_{H,j}^{(t)}\Big). \tag{8}$$

Here, the top-down feedback pathway multiplicatively gates the bottom-up signals. As a result, larger (smaller) magnitudes in the memory pathway can amplify (suppress) the corresponding magnitudes in the immediate visual pathway. This scheme allows for more direct *control* (through multiplication) of attention weights and context vectors, enabling stronger or weaker gating of specific patches.

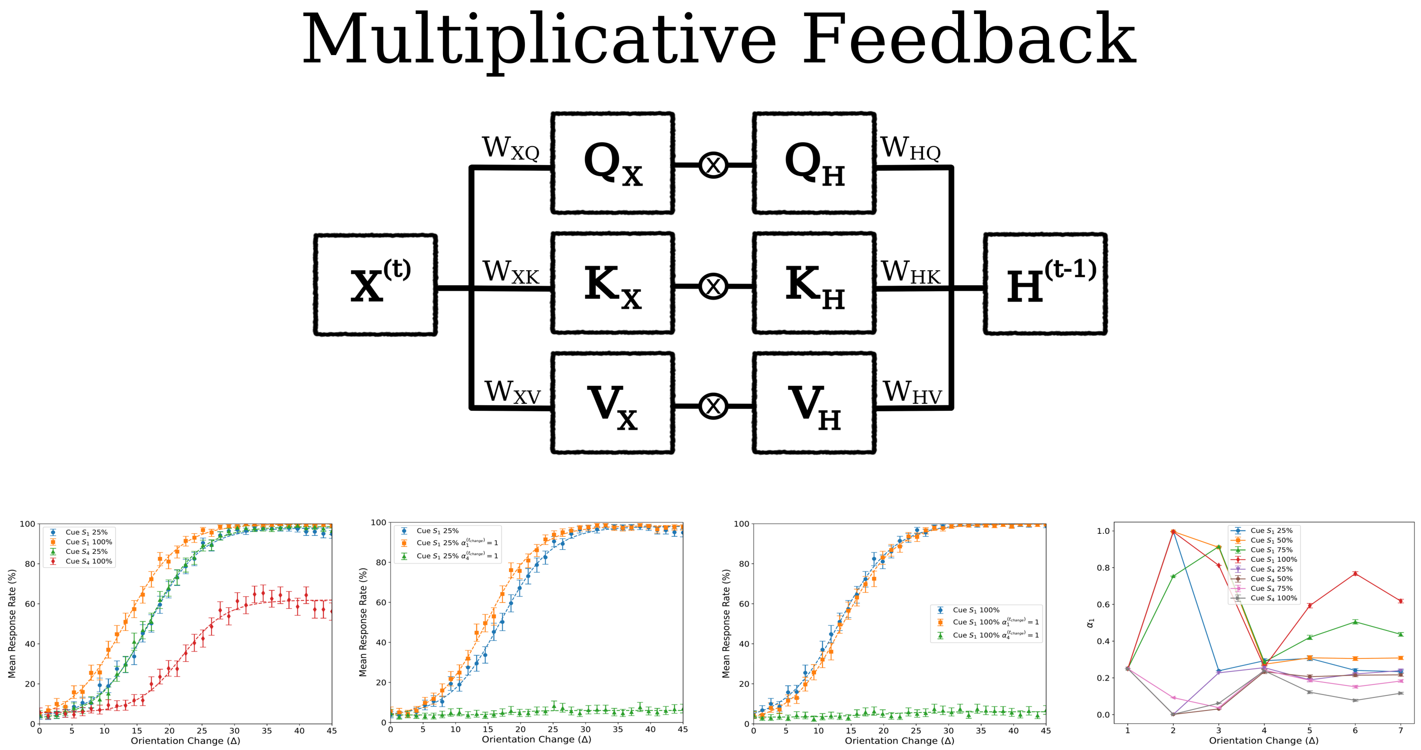

**Figure 7.** Circuit diagram and some behavioral results for a model in which recurrent feedback is implemented via the element-wise multiplication of parallel projections onto the Q, K, V self-attention components. This model shows the strongest cue modulation and the strongest attention modulation effects on behavior. In addition, the cue has a strong influence on the deployment of attention at the time of change. These factors contribute to our choice of this model as the focus of our analysis.

Additive operations can be dominated by whichever pathway has a larger magnitude, potentially diminishing subtler signals. By contrast, multiplicative modulation can act as a direct "sign-flip" mechanism or a global rescaling factor, making it inherently well-suited for precise top-down control. For instance, consider a scenario in which $\mathbf{q}_{X,i}$ or $\mathbf{k}_{X,j}$ contain elements $\pm 2$. A feedback mechanism that must flip selected signs via *addition* could require large compensatory values in $\mathbf{q}_{H,i}$ or $\mathbf{k}_{H,j}$. In contrast, a multiplicative pathway can achieve such sign flips with a scalar factor of $-1$, regardless of the original magnitude in $\mathbf{q}_{X,i}$ or $\mathbf{k}_{X,j}$.

### 2 Model Architecture

Our model integrates a Vision Transformer (ViT) with a patch-based LSTM. First, a VAE is used to pre-process the raw visual features in a purely feed-forward method. Secondly, we utilize a recurrent ViT in which self-attention has been modified to incorporate immediate and recurrent inputs in order to construct the visual percept transmitted to the patch-based LSTM. Thirdly, the LSTM utilizes the projection from the recurrent ViT to update the patch-based internal states.

#### 2.1 VAE Pre-Processing

A Variational Autoencoder (VAE) is a generative model that learns to encode input data into a latent space and reconstructs the data from this latent representation. It combines principles from deep learning and probabilistic inference, making it suitable for modeling complex data distributions. It consists of two primary components, and encoder ($F$) that encodes visual inputs to a probabilistic latent space ($z_{latent}$), and a decoder ($G$) that decodes a sampled latent vector into a visual input.

The encoder network $F$ entails multiple operations, $f \in F$ which serve to map an input image patch $\mathbf{o}_i \in \mathbb{R}^{H_{patch} \times W_{patch} \times C}$ to a latent representation characterized by a mean vector $\boldsymbol{z_\mu} \in \mathbb{R}^{d_{latent}}$ and a log-variance vector $\boldsymbol{z_{logvar}} \in \mathbb{R}^{d_{latent}}$, where $d_{latent}$ is the dimensionality of the latent space. The encoder consists of convolutional and fully connected layers as follows:

1. **First Convolutional Layer**: Applies a convolution with 16 filters, each of size $3 \times 3$, stride 2, and padding 1. This operation reduces the spatial dimensions while increasing the feature depth. The activation function is ReLU:
$$\mathbf{z_{Conv,1}} = \mathrm{ReLU}\left(f_{Conv,1}^{(1,16,3,2,1)}(\mathbf{o_i})\right)$$
2. **Second Convolutional Layer**: Applies a convolution with 32 filters, each of size $3 \times 3$, stride 2, and padding 1:
$$\mathbf{z_{Conv,2}} = \mathrm{ReLU}\left(f_{Conv,2}^{(16,32,3,2,1)}(\mathbf{z_{Conv,1}})\right)$$
3. **Flattening**: The output tensor is reshaped into a vector:
$$\mathbf{z_{flat,1}} = \mathrm{Flatten}(\mathbf{z_{Conv,2}})$$
4. **First Fully Connected Layer**: Maps the flattened vector to a 128-dimensional feature vector:
$$\mathbf{z_{flat,2}} = \mathrm{ReLU}\left(\mathbf{W}_1 \mathbf{z}_{\text{flat}} + \mathbf{b}_1\right)$$
5. **Latent Variable Parameters**: Computes the mean and log-variance vectors using two separate linear transformations:
$$\boldsymbol{z_\mu} = \mathbf{W}_\mu \mathbf{z_{flat,2}} + \mathbf{b}_\mu, \quad \boldsymbol{z_{\log \sigma^2}} = \mathbf{W}_{\log \sigma^2} \mathbf{z_{flat,2}} + \mathbf{b}_{\log \sigma^2}$$

To allow gradient-based optimization through stochastic sampling, we employ the reparameterization trick. Letting $\mu = \mathbf{z}_\mu$ and $\sigma = \exp(0.5\mathbf{z_{logvar}})$ we draw a latent vector $\mathbf{z_{latent}}$ from the approximate posterior:
$$\mathbf{z_{latent}} = \boldsymbol{\mu} + \boldsymbol{\sigma} \odot \boldsymbol{\epsilon}, \quad \boldsymbol{\epsilon} \sim \mathcal{N}(\mathbf{0}, \mathbf{I})$$
where $\boldsymbol{\sigma} = \exp\left(\frac{1}{2} \mathbf{log}\, \boldsymbol{\sigma}^2\right)$, and $\odot$ denotes element-wise multiplication.

The decoder network $G$ maps the latent vector $\mathbf{z}$ back to the reconstructed image $\hat{\mathbf{o}}_i$. The decoder mirrors the encoder but uses transposed convolutions:

1. **First Fully Connected Layer**: Transforms the latent vector to a 128-dimensional vector:
$$\hat{\mathbf{o}}_{\mathbf{flat,1}} = \mathrm{ReLU}\left(\mathbf{W}_{flat,1} \mathbf{z_{latent}} + \mathbf{b}_{flat,1}\right)$$
2. **Second Fully Connected Layer**: Maps the 128-dimensional vector to a shape suitable for convolutional layers:
$$\hat{\mathbf{o}}_{\mathbf{flat,2}} = \mathrm{ReLU}\left(\mathbf{W}_{flat,2} \hat{\mathbf{o}}_{\mathbf{flat,1}} + \mathbf{b}_{flat,2}\right)$$
3. **First Transposed Convolutional Layer**: Applies a transposed convolution with 16 filters:
$$\hat{\mathbf{o}}_{\mathbf{ConvT,1}} = \mathrm{ReLU}\left(g_{ConvT,1}^{(32,16,3,2,1,0)}(\hat{\mathbf{o}}_{\mathbf{flat,2}})\right)$$
4. **Second Transposed Convolutional Layer**: Applies a transposed convolution to reconstruct the image:
$$\hat{\mathbf{o}}_{\mathbf{ConvT,2}} = \mathrm{Sigmoid}\left(g_{ConvT,2}^{(16,1,3,2,1,0)}(\hat{\mathbf{o}}_{\mathbf{ConvT,1}})\right)$$

The VAE optimizes a loss function that combines reconstruction accuracy and the Kullback-Leibler (KL) divergence between the approximate posterior and the prior distribution. Letting $\hat{\mathbf{o}}_\mathbf{i} = \hat{\mathbf{o}}_{\mathbf{ConvT,2}}$ the loss is described as:

$$\mathcal{L} = \frac{1}{d_{image}} \|\mathbf{o}_\mathbf{i} - \hat{\mathbf{o}_\mathbf{i}}\|^2 - \beta \cdot \frac{1}{2}\left(1 + \log \boldsymbol{\sigma_i}^2 - \boldsymbol{\mu_i}^2 - \boldsymbol{\sigma_i}^2\right)$$

where $\beta$ is a hyperparameter that balances the two terms, i represents the image patch number, and $d_{image} = H_{patch} \times W_{patch} \times C$

#### 2.2 ViT

Input images $\mathbf{O}^{(t)} \in \mathbb{R}^{50\times50}$ are sub-divided into four equal patches $\{\mathbf{o_1}^{(t)}, \mathbf{o_2}^{(t)}, \mathbf{o_3}^{(t)}, \mathbf{o_4}^{(t)}\}$, with $\mathbf{o_i}^{(t)} \in \mathbb{R}^{(25\times25)}$. We found that our RL agent learned fastest, was most interpretable, and demonstrated best performance when we used the second flattend encoder layer ($\mathbf{o_{flat,2}}$) as input to the ViT (as oppose to the latent encoding). Hence, for a given patch input $\mathbf{o_i}^{(t)}$ at time $t$, we have the encoding

$$\hat{\mathbf{o}}_\mathbf{i}^{(t)} = f^*(\mathbf{o_i}^{(t)})$$

where $f^*(\cdot)$ includes encoder components (1)–(4). We also concatenate a (one-hot) positional ($\boldsymbol{\rho_i}$) and temporal ($\boldsymbol{\tau}$) encoding. Thus the full pre-processed patch input at timestep $t$ is

$$\mathbf{x_i}^{(t)} = \text{Concat}[\hat{\mathbf{o}}_\mathbf{i}^{(t)}, \boldsymbol{\rho_i}, \boldsymbol{\tau}] \tag{9}$$

The complete input to the ViT at time step $t$ is:

$$\mathbf{X}^{(t)} = (\mathbf{x_1}^{(t)}, \mathbf{x_2}^{(t)}, \mathbf{x_3}^{(t)}, \mathbf{x_4}^{(t)})^T \in \mathbb{R}^{4\times140} \tag{10}$$

The transformer computes queries, keys, and values as:

$$\mathbf{Q} = (\mathbf{X^{(t)}W_{XQ}}) \odot (\mathbf{H^{(t-1)}W_{HQ}}) \tag{11}$$
$$\mathbf{K} = (\mathbf{X^{(t)}W_{XK}}) \odot (\mathbf{H^{(t-1)}W_{HK}}) \tag{12}$$
$$\mathbf{V} = (\mathbf{X^{(t)}W_{XV}}) \odot (\mathbf{H^{(t-1)}W_{HV}}) \tag{13}$$

where $\mathbf{W_{X\cdot}} \in \mathbb{R}^{140\times140}$, $\mathbf{W_{H\cdot}} \in \mathbb{R}^{1024\times140}$, $\mathbf{H}^{(t-1)}$ is the activated memory from the previous timestep, $\odot$ denotes Hadamard product, and we have dropped the temporal superscript (implicit). Self-attention is computed as:

$$\mathbf{V_{filtered}} = \text{Softmax}(\mathbf{QK}^T)\mathbf{V} \tag{14}$$

The spatially and temporally aware visual percept is constructed as follows:

$$\mathbf{Z}^{(t)} = \mathbf{X}^{(t)} + \mathbf{V_{filtered}} \in \mathbb{R}^{4\times140} \tag{15}$$

#### 2.3 Spatial LSTM

We adapt the xLSTM architecture [3] for spatial memory. The LSTM operations are:

$$\begin{aligned}
&\tilde{\mathbf{I}}^{(t)} = \mathbf{Z}^{(t)}\mathbf{W_i} + \mathbf{H}^{(t-1)}\mathbf{R_i} && \mathbf{I}^{(t)} = \exp(\tilde{\mathbf{I}}^{(t)} - \mathbf{M}^{(t)}) && \mathbf{O}^{(t)} = \sigma(\tilde{\mathbf{O}}^{(t)})\\
&\tilde{\mathbf{F}}^{(t)} = \mathbf{Z}^{(t)}\mathbf{W_f} + \mathbf{H}^{(t-1)}\mathbf{R_f} && \mathbf{F}^{(t)} = \exp(\tilde{\mathbf{F}}^{(t)} + \mathbf{M}^{(t-1)} - \mathbf{M}^{(t)}) && \mathbf{N}^{(t)} = \mathbf{F}^{(t)} \odot \mathbf{N}^{(t-1)} + \mathbf{I}^{(t)}\\
&\tilde{\mathbf{O}}^{(t)} = \mathbf{Z}^{(t)}\mathbf{W_o} + \mathbf{H}^{(t-1)}\mathbf{R_o} && \mathbf{M}^{(t)} = \max(\tilde{\mathbf{F}}^{(t)} + \mathbf{M}^{(t-1)}, \tilde{\mathbf{I}}^{(t)}) && \mathbf{U}^{(t)} = \tanh(\tilde{\mathbf{U}}^{(t)})\\
&\tilde{\mathbf{U}}^{(t)} = \mathbf{Z}^{(t)}\mathbf{W_u} + \mathbf{H}^{(t-1)}\mathbf{R_z} && \mathbf{C}^{(t)} = \mathbf{C}^{(t-1)} \odot \mathbf{F}^{(t)} + \mathbf{U}^{(t)} \odot \mathbf{I}^{(t)} && \mathbf{H}^{(t)} = \mathbf{O}^{(t)} \odot (\mathbf{C}^{(t)}/\mathbf{N}^{(t)})
\end{aligned}$$

where $\mathbf{W_x} \in \mathbb{R}^{140\times1024}$, , $\mathbf{R_x} \in \mathbb{R}^{140\times1024}$, and all other variables $\in \mathbb{R}^{4\times1024}$. As described above, we call this a patch-based LSTM because there is a hidden state for each patch of the visual scene. Importantly, within the LSTM the hidden states are updated independently. The matrices $\mathbf{Z}^{(t)}$, $\mathbf{C}^{(t)}$, $\mathbf{H}^{(t)}$, $\mathbf{M}^{(t)}$, and $\mathbf{N}^{(t)}$ are of shape $n_{patch}$ by $d$, where $d \in d_{latent}, d_{mem}$. Right multiplication by the matrices $\mathbf{W_x}$ or $\mathbf{R_x}$ projects the latent embedding or hidden state of a specific patch to another space, independent of the other patches. By constructions, self-attention is the only mechanism by which information from visual patches (or mnemonic patches) is communicated to other patches.

#### 2.4 Actor-Critic Network

The mnemonic percept $\mathbf{H}^{(t)} \in \mathbb{R}^{4\times1024}$ serves as input to both actor and critic networks. The actor network is a 4-layer feed-forward neural network:

$$\mu_1 = \text{ELU}(H'W_1 + b_1) \tag{16}$$
$$\mu_2 = \text{ELU}(\mu_1 W_2 + b_2) \tag{17}$$
$$\mu_3 = \text{ELU}(\mu_2 W_3 + b_3) \tag{18}$$
$$\pi_\theta(a_t|H_t) = \text{Softmax}(\mu_3 W_{\text{out}} + b_{\text{out}}) \tag{19}$$

where $H' \in \mathbb{R}^{4096}$ is the flattened $H_t$. The network dimensions decrease from 4096 to 2, with the output representing the action distribution.

The critic network maps $(H_t, a_t)$ to a distributional Q-function:

$$a' = a_t W_a + b_a \tag{20}$$
$$q_0 = \text{Concat}[H', a'] \tag{21}$$
$$q_1 = \text{ELU}(q_0 W_1 + b_1) \tag{22}$$
$$q_2 = \text{ELU}(q_1 W_2 + b_2) \tag{23}$$
$$q_3 = \text{ELU}(q_2 W_3 + b_3) \tag{24}$$
$$p_\theta(q|H_t, a_t) = \text{Softmax}(q_3 W_{\text{out}} + b_{\text{out}}) \tag{25}$$

where $q_0 \in \mathbb{R}^{8192}$, and the output dimension is 15, representing discretized Q-values. The model is trained using a KL-regularized reinforcement learning objective:

$$L_Q(\theta) = \mathbb{E}_D[D_{\text{KL}}[\pi_{\text{imp}}, \pi_\theta|s_t, \tilde{\pi} = \pi_{\theta'}] + \beta D_{\text{KL}}[\Gamma_{\theta'}(q|s_t, a_t), p_\theta(q|s_t, a_t)]] \tag{26}$$

Here, $\pi_{\text{imp}}$ is the improved policy given by:

$$\pi_{\text{imp}}(a_t|s_t) \propto \exp(Q_{\theta'}(s_t, a_t)/\eta)\pi_{\theta'}(a_t|s_t) \tag{27}$$

$\Gamma_{\theta'}(q|s_t, a_t)$ is the target Q-distribution computed using the distributional Bellman operator:

$$\Gamma_{\theta'}(q|s_t, a_t) = \mathbb{E}_{s_{t+1}}\mathbb{E}_{a'\sim\pi_\theta(\cdot|s_{t+1})}\mathbb{E}_{q'\sim p_\theta(\cdot|s_{t+1},a')}[\mathbf{1}_{[q-\epsilon/2,q+\epsilon/2]}(r_t + \gamma q')] \tag{28}$$

The loss function balances policy improvement (first KL term) with Q-function learning (second KL term). The hyperparameter $\beta$ controls the trade-off between these objectives. This formulation allows for offline reinforcement learning without explicit behavior cloning, relying instead on the KL-regularization to the previous policy $\pi_{\theta'}$ to stabilize learning.

#### 2.5 Reinforcement Learning

To emulate learning processes observed in non-human primates (NHPs) and humans, our model is trained using a reinforcement learning (RL) framework [25]. In this framework, the agent interacts with its environment by observing visual stimuli and taking actions to maximize the expected cumulative rewards over time. The goal of the RL framework is to enable the model to learn optimal policies that maximize future rewards based on the agent's perceptual inputs and previous experiences. The input to the RL module is the mnemonic percept $H^{(t)}$, which encapsulates the relevant features of the visual scene as represented in the working memory module. This activated memory serves as the input to both the action-selection policy $\pi(H^{(t)})$ and the value function $V_\pi(H^{(t)})$, both of which are parameterized by neural networks in our model.

The action-selection policy $\pi(H^{(t)})$ maps the current activated memory to an action that the agent will take at time $t$. In our environment, the agent has two possible actions:

$$\pi(H^{(t)}) = 0 \quad \text{(“wait” action)},$$
$$\pi(H^{(t)}) = 1 \quad \text{(“declare change” action)}.$$

The “wait” action implies that the agent decides not to make a response and continues to process further information from the environment. The “declare change” action represents the agent's decision to identify a change in the visual stimulus. The policy network is trained to maximize the expected future rewards by choosing the action that is predicted to have the highest value. The value function $V_\pi(H^{(t)})$ estimates the expected cumulative future reward from the current activated memory $H^{(t)}$:

$$V(H^{(t)}) = \mathbb{E}\left[\sum_{\tau=t}^{T} \gamma^{\tau-t} r_\tau \,\middle|\, H^{(t)}\right], \tag{29}$$

where $\gamma \in [0, 1]$ is the discount factor, $r_\tau$ is the reward received at time step $\tau$, and $T$ is the terminal time step for the task. The value function predicts how much reward the agent expects to receive by following its learned policy $\pi$ from the current activated memory.

Learning in the RL module is driven by the temporal difference (TD) error, which measures the difference between the predicted value and the actual reward received at each time step:

$$\delta_t = r_t + \gamma V(H^{(t+1)}) - V(H^{(t)}), \tag{30}$$

where $r_t$ is the reward received at time $t$ and $\delta_t$ is the TD error. This error signal is used to update the value function $V_\pi(H^{(t)})$ and the action-selection policy $\pi(H^{(t)})$ to better predict future rewards and make more optimal decisions. In our model, both the value function and the policy are parameterized by neural networks. The agent's performance improves over time as it receives rewards and updates its predictions based on experience, thereby learning to allocate bias and make decisions that maximize cumulative rewards.

#### 2.6 Task Difficulty

To control task difficulty, Gabor stimuli were corrupted with rotational ”noise”. Defnining $\theta_i^*$ as the ”true” Gabor orientation for $S_i$, the orientation in the input image shown to the agent is:

$$\theta_i = \theta_i^* + \delta_{it}$$

where $\delta_{it} \sim \mathcal{N}(0, \sigma)$ is the rotational noise at time step $t$. If the stimulus is selected for change, then at $t = 5$ and $t = 6$:

$$\theta_i = \theta_i^* + \Delta + \delta_{it}$$

The orientation noise parameter $\sigma$ is set to 5. The orientation change parameter $\Delta$ is a random variable drawn at the beginning of a change trial, with $\Delta \sim \mathrm{U}(-k, k)$, where $k$ is adjusted based on the agent's performance, starting at $k = 65$ and decreasing as performance improves to increase task difficulty.

### 3 Logistic Function and Fitting Procedure

To model the relationship between orientation change and response rates, we employed a logistic function of the form:

$$f(x) = A + (1 - B)\frac{1}{1 + \exp(-C(x - D))} \tag{31}$$

where $x$ represents the magnitude of the orientation change, and the parameters $A, B, C, D$ govern the shape and position of the logistic curve.

- $A$ represents the lower asymptote, capturing any baseline response rate unrelated to orientation change.
- $B$ modulates the upper asymptote, accounting for deviations from a perfect detection rate.
- $C$ controls the slope of the curve, determining the rate at which response probability transitions from low to high.
- $D$ corresponds to the inflection point, the orientation change at which the response rate reaches its mid-point.

The logistic function was fitted to empirical response rate data by optimizing the parameters to minimize the discrepancy between the observed values and the model predictions. Confidence intervals for the response rates were estimated using a Bayesian credible interval approach based on Jeffreys' prior. The fitted curves provide a smooth characterization of the response behavior across different orientation change levels, allowing for a quantitative comparison across conditions.

| Cue $S_1$ | Change | A | B | C | D |
|---|---|---|---|---|---|
| 25% | $S_1$ | $0.07 \pm 0.01$ | $0.07 \pm 0.01$ | $0.30 \pm 0.01$ | $13.00 \pm 0.11$ |
| 50% | $S_1$ | $0.07 \pm 0.01$ | $0.07 \pm 0.01$ | $0.28 \pm 0.01$ | $11.89 \pm 0.14$ |
| 75% | $S_1$ | $0.06 \pm 0.01$ | $0.05 \pm 0.01$ | $0.30 \pm 0.01$ | $10.50 \pm 0.13$ |
| 100% | $S_1$ | $0.06 \pm 0.01$ | $0.06 \pm 0.01$ | $0.30 \pm 0.01$ | $9.83 \pm 0.13$ |
| 25% | $S_4$ | $0.08 \pm 0.01$ | $0.07 \pm 0.01$ | $0.27 \pm 0.01$ | $12.66 \pm 0.10$ |
| 50% | $S_4$ | $0.08 \pm 0.01$ | $0.08 \pm 0.01$ | $0.28 \pm 0.01$ | $13.18 \pm 0.12$ |
| 75% | $S_4$ | $0.09 \pm 0.01$ | $0.09 \pm 0.01$ | $0.27 \pm 0.01$ | $14.83 \pm 0.17$ |
| 100% | $S_4$ | $0.10 \pm 0.02$ | $0.24 \pm 0.02$ | $0.29 \pm 0.03$ | $17.49 \pm 0.34$ |

**Table 1.** Table showing logistic fit parameters for psychometric functions of the trained agent. Cues are always at the $S_1$ location. Standard error in parameter estimates are shown.

### 4 Decoding Analysis

#### 4.1 Decoder Architecture

To extract and interpret the information encoded within our patch-based LSTM model, we implemented a decoder architecture. This decoder is designed to process the output from various layers of the patch-based LSTM and produce task-relevant predictions. The architecture consists of a feed-forward neural network with the following structure:

$$\text{Decoder}(x) = f_3(f_2(f_1(x))) \tag{32}$$

| Cue $S_1$ | Change | x | A | B | C | D |
|---|---|---|---|---|---|---|
| 25% | $S_1$ | 1 | $0.06 \pm 0.01$ | $0.05 \pm 0.01$ | $0.29 \pm 0.01$ | $10.69 \pm 0.15$ |
| 25% | $S_1$ | 4 | $0.12 \pm 0.02$ | $0.91 \pm 0.02$ | $0.20 \pm 0.09$ | $15.64 \pm 2.63$ |
| 100% | $S_1$ | 1 | $0.07 \pm 0.01$ | $0.07 \pm 0.01$ | $0.28 \pm 0.01$ | $9.15 \pm 0.13$ |
| 100% | $S_1$ | 4 | $0.06 \pm 0.01$ | $0.92 \pm 0.01$ | $0.28 \pm 0.10$ | $14.89 \pm 1.41$ |
| 100% | $S_4$ | 1 | $0.12 \pm 0.04$ | $0.07 \pm 0.08$ | $0.27 \pm 0.07$ | $12.66 \pm 5.41$ |
| 100% | $S_4$ | 4 | $0.05 \pm 0.01$ | $0.06 \pm 0.02$ | $0.27 \pm 0.01$ | $13.42 \pm 0.18$ |

**Table 2.** Table showing logistic fit parameters for psychometric functions where we have artificially increased self-attention scores $\alpha_x^{(t_{change})}$ on either $S_1$ ($x = 1$) or $S_4$ ($x = 4$) at the time of change ($t_{change}$). Cues are at the $S_1$ location. Standard error in parameter estimates are shown.

where $x \in \mathbb{R}^{d_{\text{in}}}$ is the input vector (typically a flattened output from the patch-based LSTM), and $f_1$, $f_2$, and $f_3$ are layer functions defined as:

$$f_1(x) = \text{ELU}(\text{LN}_1(W_1 x + b_1)) \tag{33}$$

$$f_2(x) = \text{ELU}(\text{LN}_2(W_2 x + b_2)) \tag{34}$$

$$f_3(x) = W_3 x + b_3 \tag{35}$$

Here, $W_1 \in \mathbb{R}^{512 \times d_{\text{in}}}$, $W_2 \in \mathbb{R}^{256 \times 512}$, and $W_3 \in \mathbb{R}^{d_{\text{out}} \times 256}$ are weight matrices, $b_1$, $b_2$, and $b_3$ are bias vectors, $\text{LN}_1$ and $\text{LN}_2$ are layer normalization operations, and ELU is the Exponential Linear Unit activation function. The final output of the decoder is a vector in $\mathbb{R}^{d_{\text{out}}}$, with $d_{\text{out}}$ depending on the specific decoding task.

#### 4.2 Decoding Analysis of patch-based LSTM Components

##### 4.2.1 Decoding from the Complete mnemonic percept

The mnemonic percept in our patch-based LSTM, denoted as $H^{(t)} \in \mathbb{R}^{4 \times 1024}$, comprises four slots corresponding to the four image patches in our visual field. To analyze the information content of this hidden state, we flatten $H^{(t)}$ to $H_{\text{flat}} \in \mathbb{R}^{4096}$ and train a decoder to predict the location of the orientation change ($S_1$, $S_2$, $S_3$, or $S_4$). Figure 8 presents confusion matrices for our decoder evaluated on a test set. The decoder was trained on data (the mnemonic percept $H^{(t)}$ at time $t$) generated from an environment identical to that of the RL agent's training, without artificial modulations. The results in Figure 8 demonstrate that the decoder can detect change locations, albeit with suboptimal accuracy. We observe several key phenomena. First, there is a temporal enhancement effect: increased time ($t = 5$ to $t = 6$) enhances change decodability from $H$ collected at later time points, suggesting a temporal integration of change information. Secondly, inhibiting attention at a stimulus location had the effect of reducing change decodability of said stimulus, but slightly increased decodability of other change locations ($S_1$ in Figure 8, third column). Conversely, we found that directing attention towards a specific stimulus patch ($S_1$ in Figure 8, right column) slightly improves decodability for changes in $S_1$ while marginally decreasing decodability for other patches, indicating a trade-off in representational capacity.

To further investigate the impact of attentional modulation, we conducted an experiment with a forced 25% cue at the $S_1$ location, indicating a 0.25 probability of a stimulus becoming the change stimulus, given a change trial. Figure 9 presents the results of this analysis. In Figure 9, we observe several important effects. The first column shows normal decoding performance under the 25% cue condition. In the second column, we enforced a uniform Self-Attention map $A$ ($\alpha_i = 0.25$ for all $i$), which leads to a slight bias in classifying $H$ as being derived from a no-change trial. The third column shows the effect of inhibiting attention on $S_1$ ($\alpha_1 = 0$), which results in an increased number of no-change classifications when $S_1$ is the change target. This demonstrates the importance of attention for change detection in the attended location. Finally, in the fourth column, we enforced

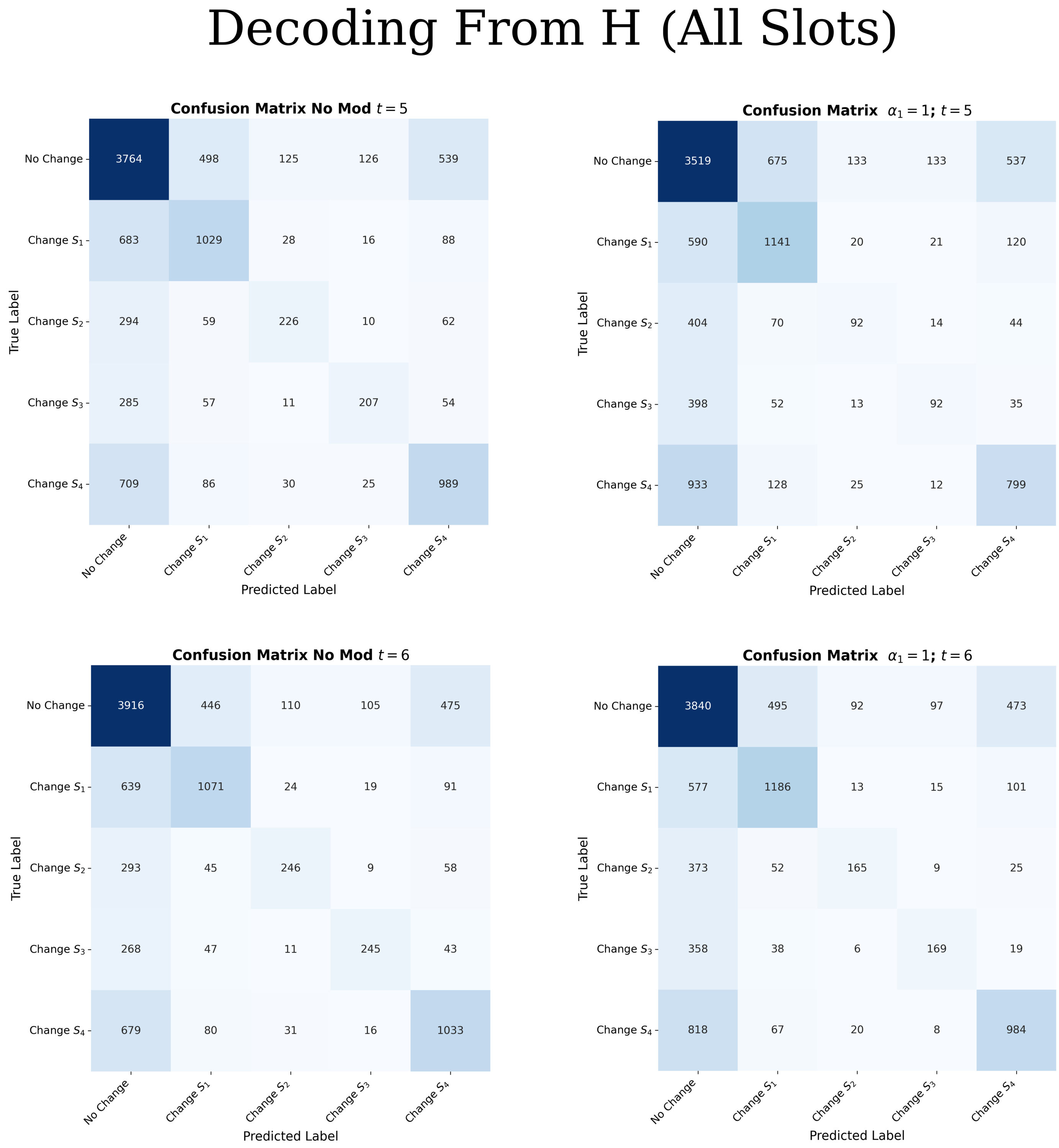

**Figure 8.** Confusion matrices for decoding change location from the mnemonic percept under various conditions.

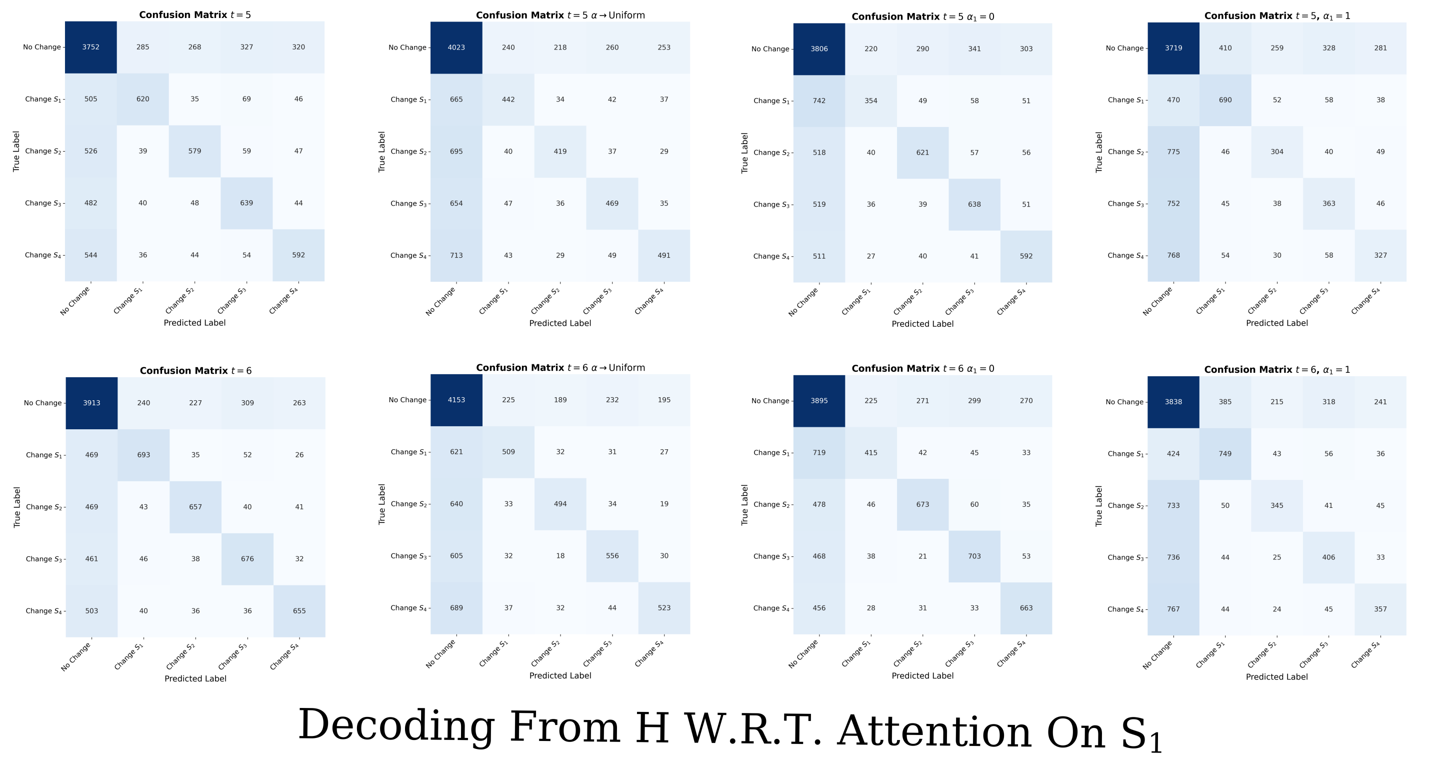

**Figure 9.** Confusion matrices for decoding change location with varying attention modulations on $S_1$.

maximum attention on $S_1$ ($\alpha_1 = 1$). This increased change classifications when $S_1$ was the change target but decreased change classification for other targets.

###### 4.2.2 Decoding from a Single Mnemonic Percept Patch

To investigate the information transfer between patches facilitated by self-attention, we decoded exclusively from the first mnemonic percept patch of $H^{(t)}$, $h_1^{(t)} \in \mathbb{R}^{1024}$, corresponding to the $S_1$ location. It is important to note that the change location was randomly selected in all trials. Figure 10 presents the results of this analysis. Our results reveal several key findings. In the case where attention is not modulated (first column), the decoder can correctly classify the occurrence of change in the majority of trials just from the $h_1^{(t)}$ patch, indicating that a single mnemonic patch contains information about changes across all patches (the probability of a change occurring at $S_1$ is 0.2). When we force maximum attention on $S_1$ ($\alpha_1^{(t)} = 1$, second column), we observe a loss of 'change' classifications on change trials (i.e., can only reliably decode the changes from the $S_1$ location). This suggests that maximal attention on $S_1$ suppresses the propagation of change signals from other patches to the $S_1$ slot. When we force no attention on $S_1$ ($\alpha_1 = 0$, not shown), the decodability of change from slot $S_1$ is only slightly affected. This is because $S_1$ changes can be decoded without attention due to direct access to $h_1^{(t)}$ patch information. These results demonstrate the crucial role of self-attention in propagating change information across memory slots.

###### 4.2.3 Decoding from the Actor Network's First Activation Layer

To understand how the actor network processes the information from the patch-based LSTM, we trained a decoder on the first activation layer of the actor network. Figure 11 presents the results of this analysis. The results in Figure 11 reveal several important aspects of information processing in the actor network. The left two columns show that much of the spatial information is lost in this first activation layer, as evidenced by the poor classification of orientation change locations compared to decoding from the hidden state (Figure 8). This

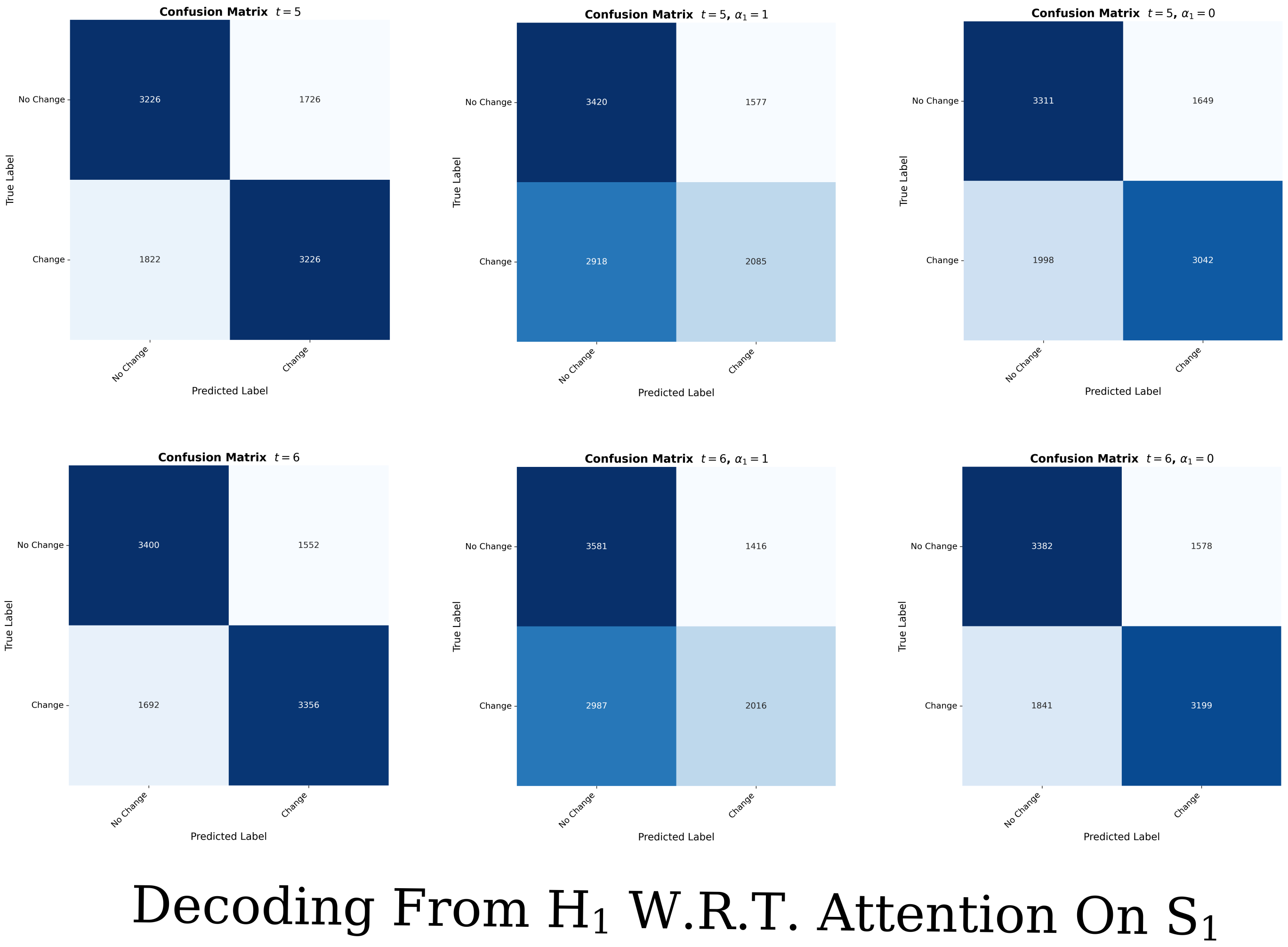

**Figure 10.** Confusion matrices for decoding change occurrence from the first mnemonic percept patch under varying attention modulations.

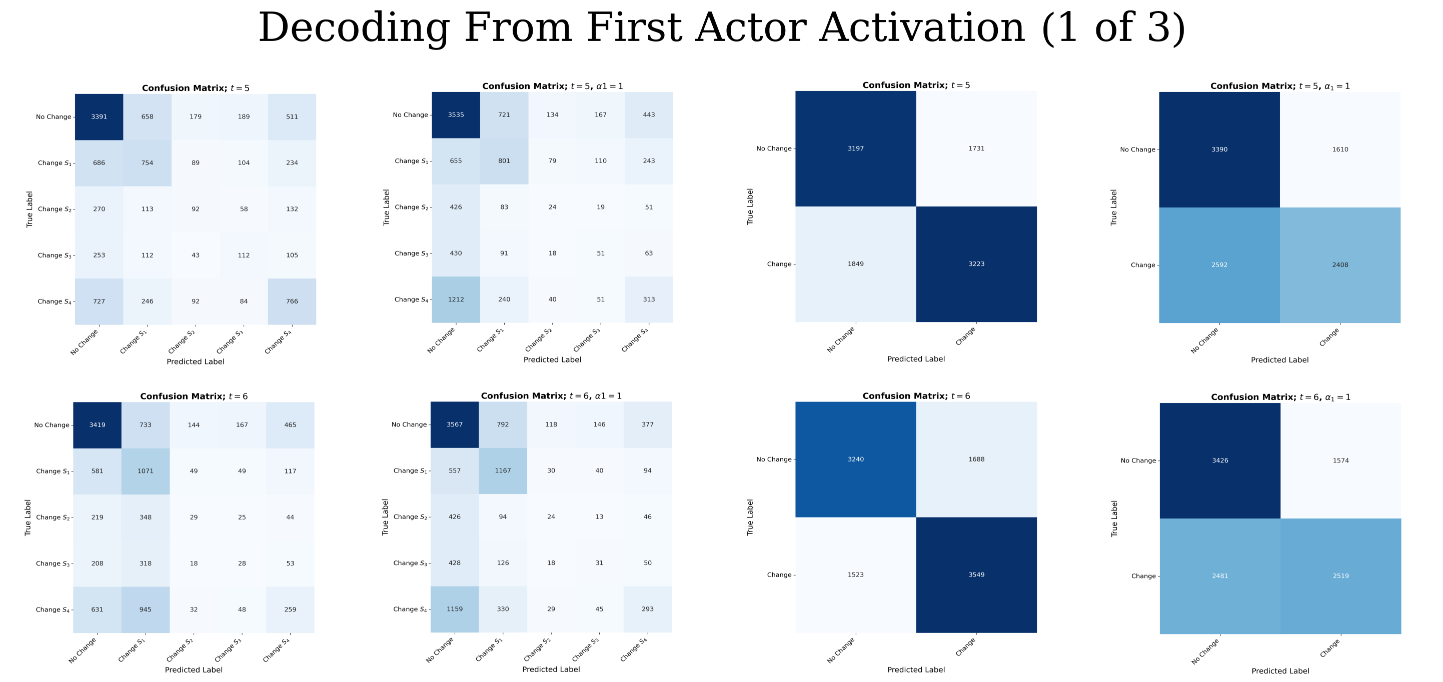

**Figure 11.** Confusion matrices for decoding change location and occurrence from the actor network's first activation layer.

suggests a loss of spatial specificity in the actor network. However, the right two columns demonstrate that changes can still be successfully decoded from this layer, indicating that change information is preserved and potentially emphasized in the actor network.

When we force $\alpha_1 = 1$ (rightmost column), we observe a loss of change classification. Importantly, this is not due to the absence of the change signal (as it is still present in the memory slot associated with the change location) but rather due to a weaker presence of the change signal. This suggests that self-attention serves to amplify the change signal by propagating it across all memory slots.

To further investigate the impact of attentional modulation on change detection in the actor network, we conducted an additional experiment focusing only on $S_1$ changes with varying levels of attention inhibition. Figure 12 presents these results. In Figure 12, we observe a clear progression as the inhibition of attention is relaxed. When self-attention on $S_1$ is completely inhibited (left column), the model fails to classify the majority of changes. However, the change signal is not entirely eradicated, as evidenced by the non-zero number of change classifications. As inhibition decreases (right columns), the confusion matrices show increasingly accurate classification structures. This demonstrates the graded nature of attentional modulation in the detection of changes in the actor network.

###### 4.2.4 Analysis of Actor Network Logits

To gain insight into the decision-making process of the actor network, we analyzed the logits of its final layer (pre-softmax activation). Figure 13 presents this analysis. Figure 13 reveals several key aspects of the decision-making process of the actor network. First, we observe a clear inverse linear relationship between 'Declare Change' and 'Wait' logits, indicating a competitive decision-making process. Second, we note a strong dependence on change intensity: 'Wait' logits tend to be high when change intensity ($\Delta$) is low, while 'Declare Change' logits are high for large $\Delta$ values. This suggests that the actor network has learned to respond in a graded manner with respect to the change signal. Interestingly, the spatial location of the change does not show any obvious influence on this relationship, indicating that the actor network has learned to make decisions based primarily on change signal independent of change location.

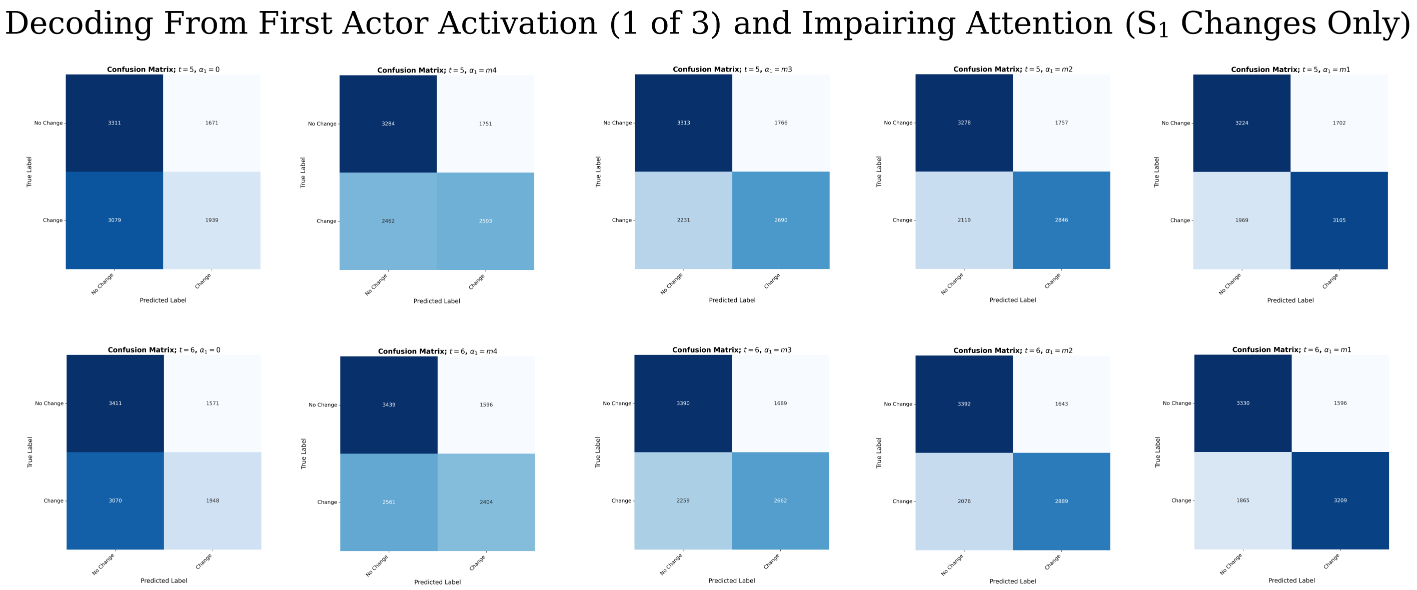

**Figure 12.** Confusion matrices for decoding $S_1$ changes from the actor network's first activation layer with varying levels of attention inhibition.

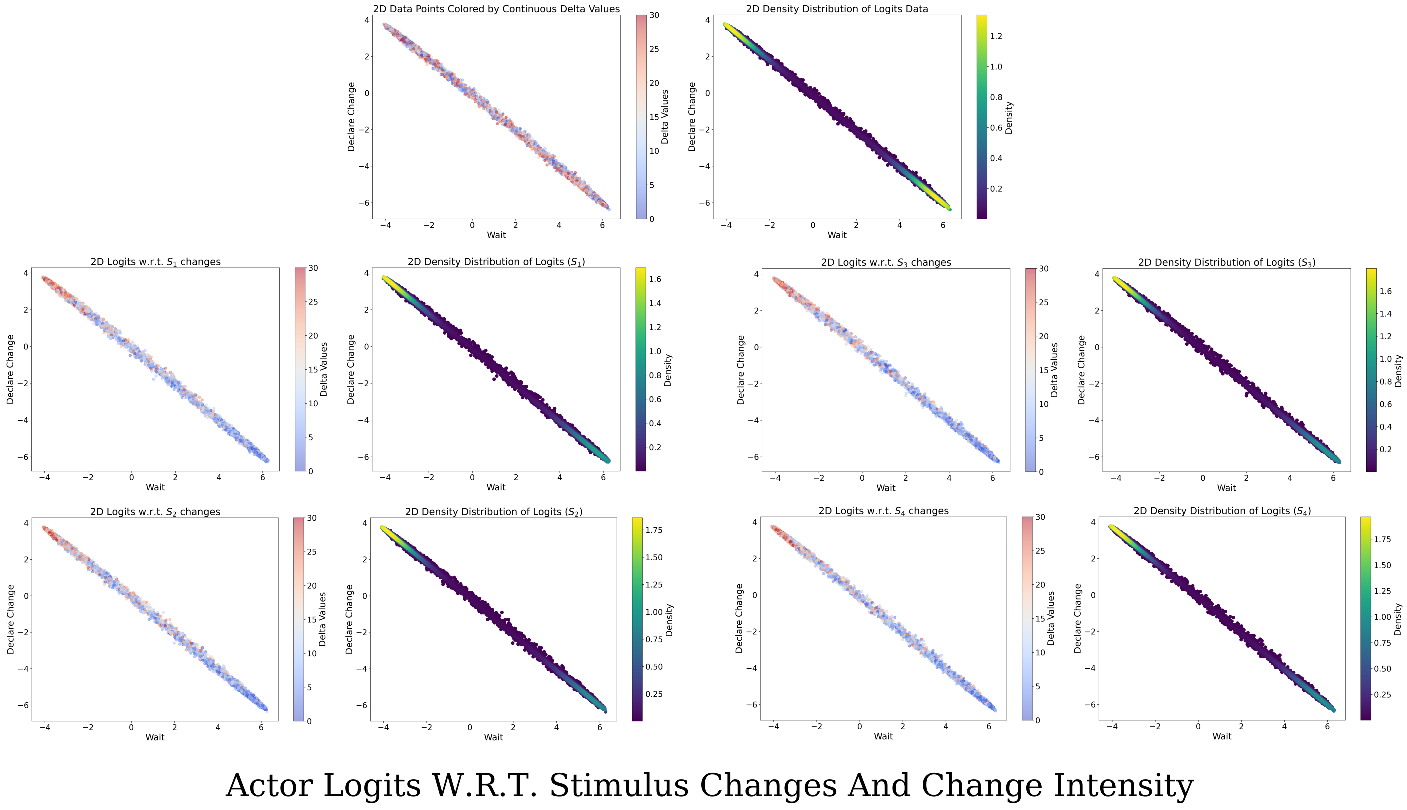

**Figure 13.** Relationship between 'Declare Change' and 'Wait' logits in the actor network's final layer.

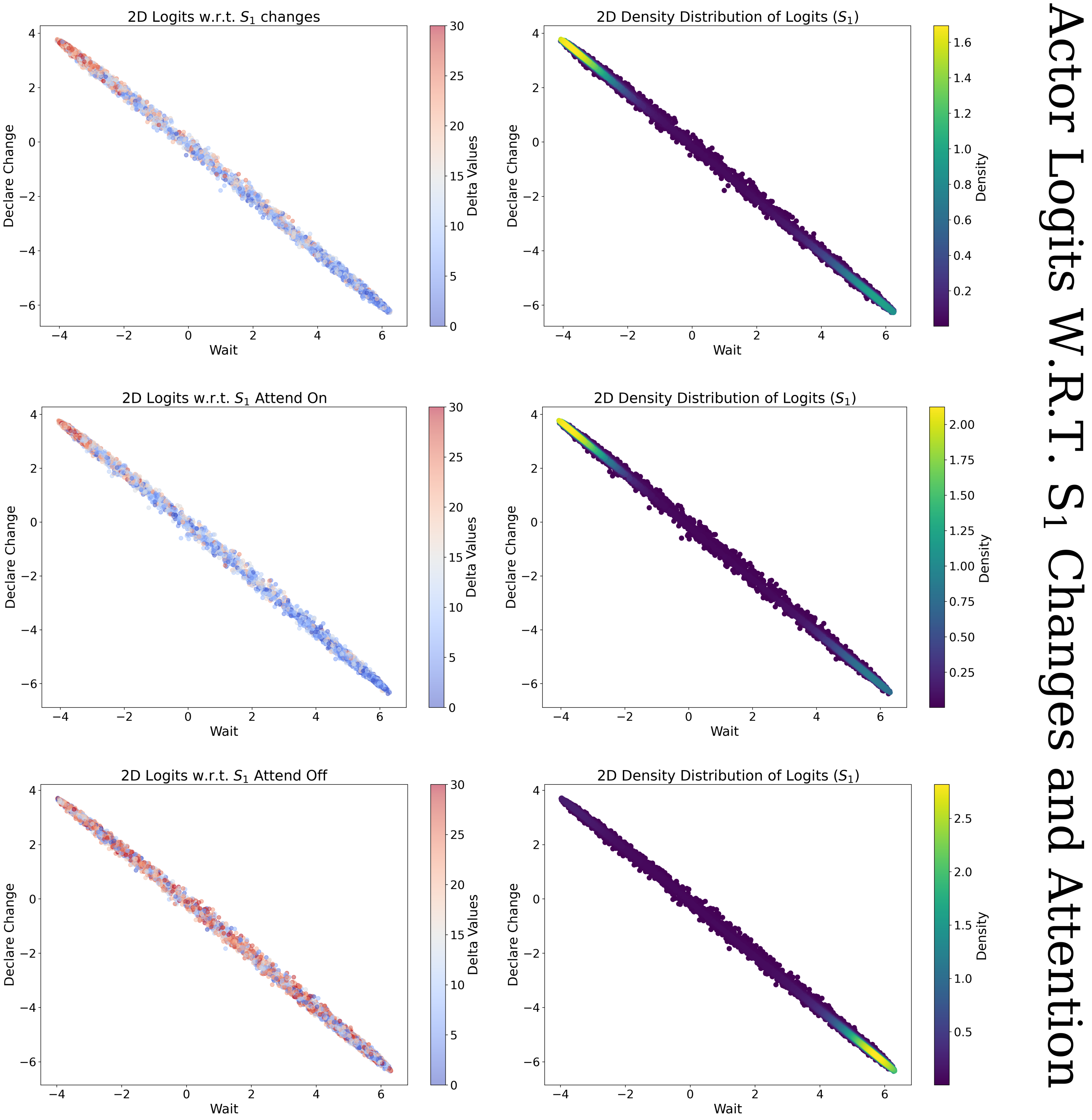

**Figure 14.** Effect of attentional modulation on actor network logits for $S_1$ changes.

To investigate the impact of attentional modulation on the actor network's decision-making process, we analyzed the logit structure under varying attention modulations for $S_1$ changes. Figure 14 presents these results. The results in Figure 14 demonstrate the significant impact of attentional modulation on the decision-making process of the actor network. Maximizing $\alpha_1$ slightly increases the density of points in the high 'Declare Change' logit space, suggesting that increased attention facilitates change detection. Conversely, minimizing $\alpha_1$ significantly decreases the density of logits in the high-valued 'Declare Change 'logit space and increases the density in the high-valued 'Wait 'logit space. This demonstrates that attentional withdrawal substantially impairs the network's ability to detect changes.

##### 4.3 Self-Attention Amplifies Change Signals

Let $C^{(t)} \in \mathbb{R}^{4\times 1024}$ represent the mnemonic state of the Recurrent ViT at time $t$, where each row corresponds to a memory slot associated with one of the four visual patches. Additionally, let $k^*$ be the indexed location of the stimuli that experienced an orientation change, $S_{k^*}$, and $\psi(\cdot)$ be some measure of the strength of a change signal. While the many parameters and complexity make a closed form description of how a change signal is communicated among memory slots challenging, based on the above results and structure of the Recurrent ViT, we can characterize some key properties of the process. From the structure of the Recurrent ViT, we know that self-attention is the only mechanism that allows information to flow from one mnemonic patch to another. Thus, if $i \neq k^*$ and $a^{(t)}_{i,k^*} = 0 : \forall t \geq t_{change}$, then $\psi(c^{(t)}_i) = 0$, where $c^{(t)}_i$ is the memory patch associated with patch $i$. Additionally, we know that if $i = k^*$, then attention is not needed to propagate change information to the memory patch $c^{(t)}_i$, i.e., $\psi(c^{(t)}_i; a^{(t)}_{i,k^*}) \approx \psi(c^{(t)}_i; 0) : \forall t \geq t_{change}$. Finally, we know that if we apply this measure to all memory patches, $\psi(C^{(t)})$, we know that $\psi(C^{(t)}; \alpha_{k^*})$ increases as $\alpha_{k^*}$ increases, where

$$\alpha_{k^*} = \sum_{i=1}^{4} a_{i,k^*}$$

Since the first layer of the actor network seems to corrupt spatial information but preserve binary change information Figure 11, we can extrapolate that the primary function of self-attention in this task is to amplify change signals.

### 5 Influence of Induced Bias on Value Estimates and Temporal Difference Errors

Reinforcement learning (RL) provides a computational framework for modeling how agents learn to make decisions through interactions with their environment [25]. In neuroscience, RL has been utilized to explain how animals and humans adapt their behavior based on reward feedback [5, 21]. A key component of RL is the *value function* $V(H^{(t)})$ (Equation 29), which estimates the expected cumulative future reward from a given the mnemonic percept $H^{(t)}$ at time $t$. The value function is updated using the *temporal difference (TD) error* $\delta_t$ (Equation 30), which quantifies the discrepancy between expected and received rewards. In the brain, dopaminergic neurons have been associated with encoding TD errors [12, 24]. Fluctuations in dopamine release have been found to correspond to reward prediction errors, influencing synaptic plasticity and learning processes [8]. Attention mechanisms, which prioritize certain sensory inputs over others, could be motivated by value [1, 2, 7, 10, 18] while also potentially influencing value estimates [15, 16, 19]. Biasing attention toward specific stimuli may thus modulate the computation of TD errors.

##### 5.1 Effects of Bias Manipulation on Temporal Difference Errors

In Figure 15, we present the TD errors and value estimates from our model under various conditions. Figure 15A compares the TD errors when a cue is presented at location $S_1$, and a change occurs either at $S_1$ or $S_4$. When a high-validity cue is shown at a location different from where the change occurs (dashed red line in Figure 15A), we observe significant suppression of the TD errors. Figures 15B and 15C demonstrate the influence of bias manipulations on the TD errors. For low-validity cues, maximizing the bias on the change location (setting $\alpha_1^{(t_{change})} = 1$: red curves in Figure 15B and C) results in significant increases in the TD error (red line in Figure 15B). However, for high-validity cues, even a maximized bias has negligible effects (overlapping curves in Figure 15C). When we force a high bias at locations other than the change location (Figures 15E and 15F), we observe a nearly complete suppression of the TD errors. In summary, the TD errors significantly increase with respect to the bias when the biased location and the change location coincide (red, blue, and green dashed lined in Figure 15D). However, when the change location and the assigned bias do not coincide, or a high validity cue was shown at a location different than the change location, we see a significant suppression of TD errors (dashed blue, dashed red, and dashed purple lines in Figure 15D).

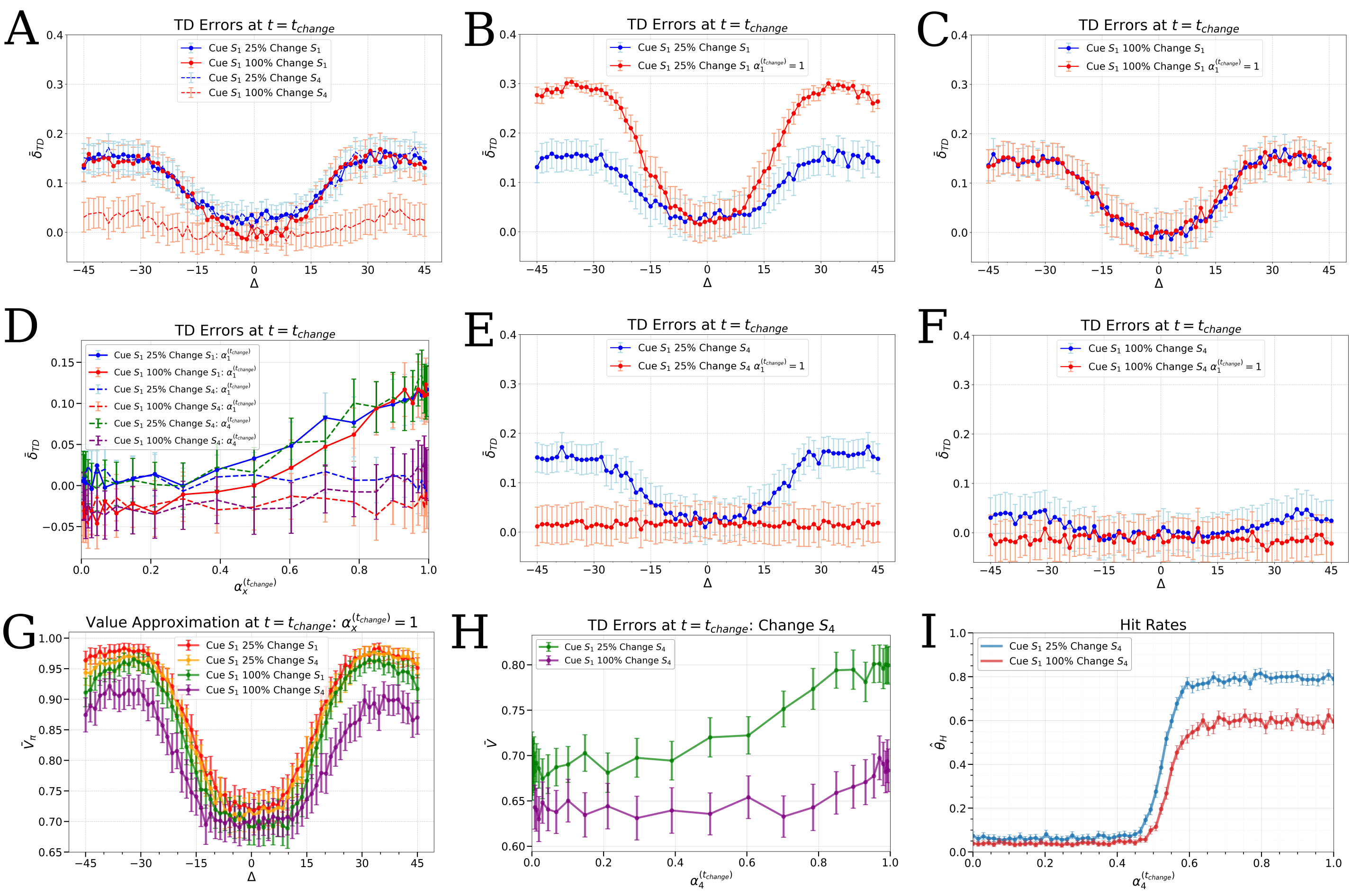

**Figure 15.** Plots showing the TD errors ($\delta_{TD}$ from Equation 30 in **A–F**) and value estimates ($V_\pi$ from Equation 29 in **G** and **H**). Data points are averaged over 500 trials. Cues always occur at the $S_1$ location and changes occur at either the $S_1$ or $S_4$ location. In **D**, bias is modulated with respect to the bias indicated in the legend, i.e., $\alpha_x^{(t_{change})} = \alpha_4^{(t_{change})}$ for the dashed purple line. In **G**, bias is maximized with respect to the change location for all $\Delta$ shown, i.e., for the purple curve $\alpha_4^{(t_{change})} = 1$. In **I** the hit-rate is shown for modulation of bias $\alpha_4^{(t_{change}}$ and changes on $S_4$

#### 5.2 Effects of Bias Manipulation on Value Estimates

The cue location, probability assignment by the cue, and the induced bias all affect the TD errors at the time of change. Similar to the method used to generate Figures 16H and 16K, we sought to isolate the effects of the cue from the secondary bias manipulation the cue induces at the time of change. Therefore, in Figure 15G, we varied the cue and change conditions but maximized the bias on the change location at the time of change (setting $\alpha_1^{(t_{\text{change}})} = 1$ for a change occurring at location $S_1$). A high-validity cue on a location different from the change location causes a decrease in value estimates, even for large orientation changes $\Delta$ (purple lines in Figure 15G). We also show that the value estimate increases approximately linearly with the bias, $\alpha_4^{(t_{change})}$, when there is a low-validity cue at an opposing location and the change location ($S_4$) differs from the cued location (green line in Figure 15H). The value estimation is significantly suppressed when there is a high-validity cue on a location different from the change (purple line in Figure 15H).

Considering the results shown in Figures 15G and 15H together with those in Figure 16H, we might be tempted to attribute the increased decision criterion resulting from high validity cues to a decrease in perceived value, since our model's action selection is based solely on value estimates. However, in Figure 15I, we demonstrate that there is a sharp increase in response rates before any such increase in value estimates (red curve in Figure 15I compared to the purple curve in Figure 15H). This suggests that there is some level of generalization beyond that of value estimates in our model's action selection policy.

### 6 Manipulating Bias Influences Criterion and Sensitivity

#### 6.1 Criterion and Sensitivity in Signal Detection Theory

Signal detection theory provides a framework for analyzing how observers detect signals in the presence of noise. Two key measures in this theory are criterion and sensitivity. The criterion represents the observer's decision threshold for reporting a signal as present. Mathematically, it is defined as:

$$c = -\frac{1}{2}(z(\theta_H) + z(\theta_{FA}))$$

Where $z(\theta_H)$ is the z-score of the hit rate ($\theta_H$) and $z(\theta_{FA})$ is the z-score of the false alarm rate ($\theta_{FA}$). Intuitively, the criterion reflects how liberal or conservative the observer is in reporting signals. A lower (more negative) criterion indicates a liberal bias - the observer is more willing to report signals even with weak evidence. A higher (more positive) criterion indicates a conservative bias - the observer requires stronger evidence to report a signal.

Sensitivity, often denoted as $d'$ (d-prime), measures the observer's ability to discriminate between signal and noise. It is calculated as:

$$d' = z(\theta_H) - z(\theta_{FA})$$

Sensitivity represents the standardized difference between the means of the signal and noise distributions. A higher $d'$ indicates better ability to distinguish signal from noise. These measures allow researchers to separate an observer's inherent sensitivity to signals from their decision-making strategy (criterion). By analyzing both, we can understand not just how well an observer detects signals, but also their underlying decision-making processes.

#### 6.2 Random Variable Interpretation of the Hit-Rate and False-Alarm Rate

Let $X_H$ and $X_{FA}$ be random variables associated with the agent's outcome on a single trial. On change trials, when $X_H = 1$, the agent successfully detected a change and the outcome is a hit. If $X_H = 0$, the agent did not detect a change and the outcome of the trial is a miss. On no-change trials, $X_{FA} = 1$ if the outcome of the trial was a false alarm and $X_{FA} = 0$ when the outcome is a correct reject. If we condition on the trial type, then the two random variables are independent, i.e.,

$$X_H|\{\text{“Change Trial”}\} \perp X_{FA}|\{\text{“No-Change Trial”}\}$$

If we let $\theta_H$ and $\theta_{FA}$ be the hit-rate and false-alarm-rate, then

$$\hat{\theta}_H = \frac{n_H}{n_H + n_M} \quad \hat{\theta}_{FA} = \frac{n_{FA}}{n_{FA} + n_{CR}} \tag{36}$$

where $\hat{\theta}_H$ and $\hat{\theta}_{FA}$ is an estimator for the true hit-rate and false-alarm-rate, $\theta_H$ and $\theta_{FA}$. For brevity, let $X_p$ and $\theta_p$ be such that $p \in$ {H,FA}, and assume the trial conditioning described above. Given a sequence of $n$ random variables, $\{X_p^{(i)}\}_{i=1}^n$, we can model $\theta_p$ as

$$\hat{\theta}_p = \frac{1}{n}\sum_{i=1}^{n} X_p^{(i)}$$
$$X_p^{(i)} \sim \text{Bern}(\theta_p)$$

where Bern$(\theta_p)$ is a Bernoulli random variable with parameter $\theta_p$. By the law of large numbers, we know that this estimator converges to the true parameter in the large sample limit

$$\lim_{n\to\infty} \frac{1}{n}\sum_{i=1}^{n} X_p^{(i)} = \theta_p$$

Using the Central Limit Theorem (CLT), we can also compute the variance of this estimate. By CLT,

$$\sqrt{n}\left[\hat{\theta}_p - \theta_p\right] \to \mathcal{N}(0, \hat{\sigma}_p^2)$$

where $\hat{\sigma}_p^2 = \hat{\theta}_p(1-\hat{\theta}_p)$ and $\hat{\sigma}_p^2 \to \sigma_p^2$ in the large sample limit. Thus, the variance of our parameter estimate is

$$Var(\hat{\theta}_p) \approx \frac{\hat{\sigma}_p^2}{n}$$

#### 6.3 z-Score as a Random Variable Transformation

If we treat $\hat{\theta}_p$ as a random variable, then computing the z-score is a transformation of this random variable. Hence, we can use the Delta Method to derive the approximate properties of this transformation. By the Delta Method:

$$\sqrt{n}\left[z(\hat{\theta}_p) - z(\theta_p)\right] \to \mathcal{N}\left(0, \hat{\sigma}_p^2[z'(\hat{\theta}_p)]^2\right)$$

where $z'(\hat{\theta}_p) = \frac{dz}{dx}|_{\hat{\theta}_p}$. This gives

$$Var(\hat{\theta}_p) \approx \frac{\hat{\sigma}_p^2}{n} z'(\hat{\theta}_p)$$

We can evaluate $z'(\hat{\theta}_p)$ by using the Inverse Function Theorem (IFT). Let $\psi(x)$ be the standard normal probability density function, i.e.,

$$\psi(x) = \frac{1}{\sqrt{2\pi}} X^{\frac{-x^2}{2}}$$

Then the cumulative distribution function is

$$\phi(x) = \int_{-\infty}^{x} \psi(t)dt$$

This gives $z'(\hat{\theta}_p) = \frac{d\phi^{-1}}{dx}|_{(\hat{\theta}_p)}$. By IFT,

$$\begin{aligned} z'(\hat{\theta}_p) &= \left.\frac{d\phi^{-1}}{dx}\right|_{\hat{\theta}_p} \\ &= \frac{1}{\left(\frac{d\phi}{dx}\big|_{\phi^{-1}(\hat{\theta}_p)}\right)} \\ &= \frac{1}{\psi(\phi^{-1}(\hat{\theta}_p))} \end{aligned}$$

Together with the above, this yields

$$Var(\hat{\theta}_p) \approx \left(\frac{\hat{\sigma}_p^2}{n}\right)\left(\frac{1}{\psi(\phi^{-1}(\hat{\theta}_p))}\right)$$

#### 6.4 Criterion and Sensitivity are Normally Distributed in the Large Sample Limit

Since we are conditioning on the trial type, our estimates of the hit-rate and false-alarm-rate are independent. Therefore, the criterion and sensitivity consists of the summation of two independent normal (approximate) random variables. For the estimate ($\hat{c}$) of a true criterion ($c$), we have

$$\hat{c} \sim \mathcal{N}\left(c, \frac{1}{4}\left(\frac{\hat{\sigma}_H^2}{n_{CT}}\left(\frac{1}{\psi(\phi^{-1}(\hat{\theta}_H))}\right) + \frac{\hat{\sigma}_{FA}^2}{n_{NT}}\left(\frac{1}{\psi(\phi^{-1}(\hat{\theta}_{FA}))}\right)\right)\right)$$

For the estimate ($\hat{d'}$) of a true sensitivity ($d'$), we have

$$\hat{d'} \sim \mathcal{N}\left(d', \left(\frac{\hat{\sigma}_H^2}{n_{CT}}\left(\frac{1}{\psi(\phi^{-1}(\hat{\theta}_H))}\right) + \frac{\hat{\sigma}_{FA}^2}{n_{NT}}\left(\frac{1}{\psi(\phi^{-1}(\hat{\theta}_{FA}))}\right)\right)\right)$$

To quantify the effects of bias modulation further, we analyzed the decision criterion and sensitivity measures under different bias manipulations. In signal detection theory, an observer's ability to distinguish between signal and noise is measured using two key metrics: sensitivity ($d'$) and criterion ($c$). Sensitivity measures how well the observer can distinguish between signal and noise, calculated using the hit rate ($\hat{\theta}_H$) and the false alarm rate ($\hat{\theta}_{FA}$). The formula for $d'$ is:

$$d' = Z(\hat{\theta}_H) - Z(\hat{\theta}_{FA}), \tag{37}$$

where $Z$ represents the inverse cumulative distribution function (CDF) of the standard normal distribution, also known as the z-score (not to be confused with the visual percept $Z^{(t)}$ in our model). Criterion measures the observer's decision bias, indicating the tendency to favor one type of response over another:

$$c = -\frac{1}{2}\left[Z(\hat{\theta}_H) + Z(\hat{\theta}_{FA})\right]. \tag{38}$$

We assessed these metrics for individual stimulus locations by modifying the task environment so that only the stimulus location in question could experience an orientation change, with change trials occurring with probability $0.5$. For example, if we are interested in the criterion of the agent's response with respect to stimulus location $S_4$ given a $100\%$ validity cue at $S_1$, then we run multiple trials with the cue at $S_1$ and only allow orientation changes to occur at $S_4$. We then compute Equations (37) and (38) over the obtained behavioral data.

#### 6.5 Bias effects on criterion and sensitivity

In the literature, there is ongoing debate regarding how attention modulates criterion and sensitivity in visual tasks. Some studies have found that attention can modulate both the decision criterion and sensitivity, indicating that attention enhances perceptual processing and decision-making [17]. Specifically, Luo and Maunsell [17] showed that attentional cues not only improve the ability to discriminate between stimuli (increasing sensitivity) but also affect the observer's response bias (criterion). Contrastingly, recent experimental work suggests that presaccadic attention may predominantly influence the decision criterion rather than sensitivity. For instance, Gupta et al. [9] found that shifts of attention just before eye movements adjust the criterion without significantly affecting sensitivity, implying that attention reallocations linked to saccades might alter decisional strategies more than perceptual encoding. Other studies propose that strategically modulating the decision criterion can indirectly enhance sensitivity. Wang et al. [28] argue that criterion shifts, when optimized, can lead to improvements

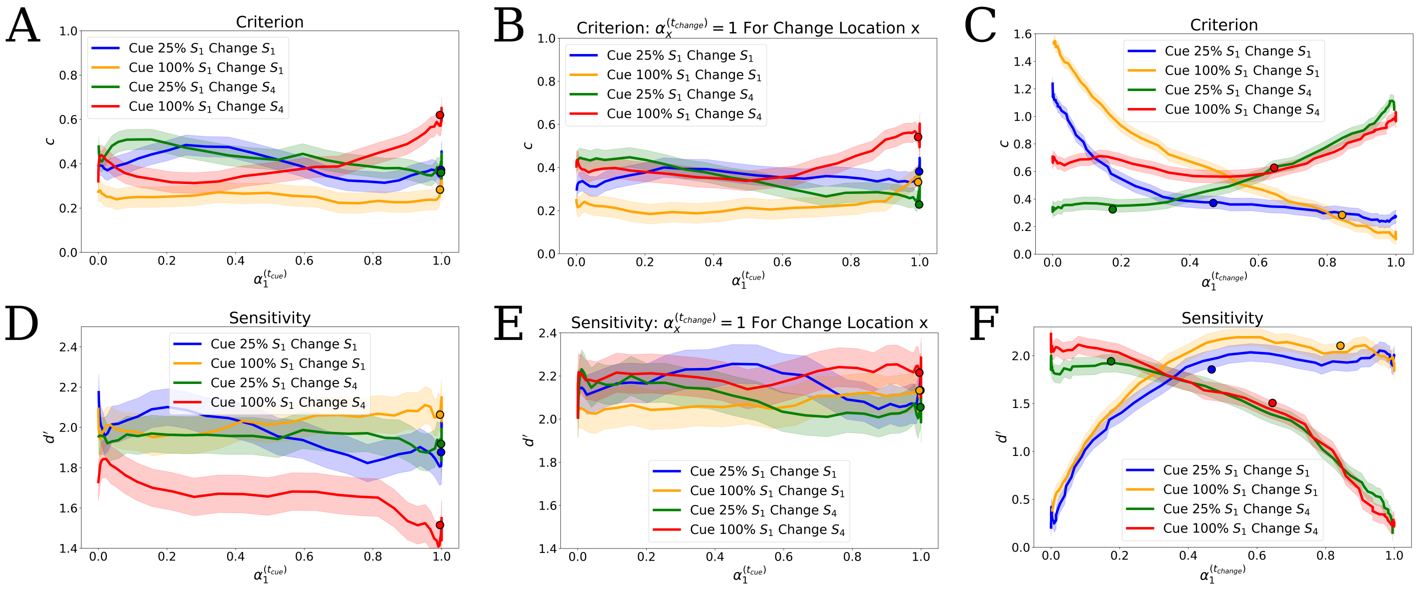

**Figure 16.** Plots showing the effect of artificially modulating the bias. All data points are the result of an average over 500 trials. Artificial modulation involves inducing a high bias in one of the columns of the self-attention map. In all cases, the bias is induced on the competitors with respect to the $S_1$ ($\alpha_1^{(t)}$ and $\xi_1^{(t)}$) location or the $S_4$ ($\alpha_4^{(t)}$ and $\xi_4^{(t)}$) location. In **A–C**, we plot the hit rates versus the orientation change $\Delta$. If $\alpha_i^{(t_{\text{change}})} = 1$, this indicates that the visual percept $Z^{(t_{\text{change}})}$ has been completely biased toward $\xi_i^{(t_{\text{change}})}$, i.e., there are no traces of other competitors in the visual percept. We manipulate the bias to this extreme for either the $\xi_1^{(t_{\text{change}})}$ or $\xi_4^{(t_{\text{change}})}$ competitors in **A–C** and then again for the reaction time plots in **D–F**. In **G–I**, we show the criterion measured (Equation 38) after manipulating the bias with respect to the $S_1$ location at the time of cue ($\alpha_1^{(t_{\text{cue}})}$ in **G** and **H**) or at the time of orientation change ($\alpha_1^{(t_{\text{change}})}$ in **I**). The colored dots indicate the average criterion measured without bias manipulations. The criterion was evaluated over 500 trials where there was a 50% chance of the change occurring at the indicated locations; otherwise, no change occurred. This procedure is repeated in **J–L** with the exception that we are measuring the sensitivity (Equation 37). In **H** and **K**, we modulate the bias with respect to the $S_1$ location at the time of cue ($\alpha_1^{(t_{\text{cue}})}$), but then immediately following the cue, we maximize the bias for the location where the change will occur ($\alpha_x^{(t_{\text{change}})}$ for a change on $S_x$, $x \in \{1, 4\}$). This is to prevent the secondary biasing effects that the cue induces at change time from affecting our comparisons.

in sensitivity by better aligning the decision boundary with the signal distribution, particularly in tasks with asymmetric signal distributions or unequal prior probabilities. This suggests a complex interplay between criterion and sensitivity, where attentional mechanisms can influence perceptual performance both directly and indirectly.

Our findings contribute to this debate by showing that artificially manipulating the bias in our model can differentially affect criterion and sensitivity. In Figure 16G and J, we manipulated the bias at the time of cue $t_{\text{cue}}$ with respect to the cue location (e.g., adjusting $\alpha_1^{(t_{\text{cue}})}$ for a cue at $S_1$). We observed that maximizing the bias at the cue location, for a high validity assigning cue, increases the criterion and decreases sensitivity when the change occurs at a different location (red curves in Figure 16G and J). This suggests that the model's attentional focus at the cue time can affect its ability to detect changes elsewhere. Notably, under natural conditions without artificial bias manipulations, attention is already maximized at the cue location (dots in Figures 16G, H, J, K).

We know that spatial bias at the time of change is affected by the attributes of the cue. For example, a 100% valid cue at the $S_1$ location at the time of cue leads to higher bias on the cued location at the time of change. To isolate the effect of bias at the time of cue from secondary effects at the time of change, we conducted additional experiments where we maximized the bias for the change location at $t_{\text{change}}$ (Figures 16H and K). This negated the secondary bias effect resulting from the cue and offers more equality in comparisons. We found that the cue's effect on the criterion remained largely unchanged (Figure 16H), but the impact on sensitivity was abolished (Figure 16K), indicating that the decrease in sensitivity observed in Figure 16J is a result of spatial bias being directed away from the change location at the time of change.

In Figures 16I and L, we demonstrate that manipulating the bias at the time of change ($\alpha_1^{(t_{\text{change}})}$) significantly influences critetion and sensitivity. Increasing bias toward the change location decreases the criterion and increases sensitivity (blue and gold curves in Figure 16I and L), while increasing bias toward non-change locations has the opposite effect (green and red curves in Figure 16I and L). This occurs because increasing $\alpha_i^{(t_{\text{change}})}$ amplifies the expression level of the spatially corresponding internal representation $\xi_i^{(t)}$. If the change occurred at stimulus location $S_i$ at time $t = t_{\text{change}}$, then the change information would initially be localized in $\xi_i^{(t_{\text{change}})}$, since $xi_i^{(t)}$ is the only competing internal representation that consists of information from the immediate visual patch $x_i^{(t)}$. Increasing $\alpha_i^{(t_{\text{change}})}$ increases the signal strength, or expression, of this change information in the visual percept $Z^{(t)}$ to working memory. Since biasing the internal representation associated with the change location is directly amplifying the most task relevant information in the visual percept, it seems more fitting to describe attention at the time of change as mostly modulating sensitivity, and that criterion decreases is a result of conversions from misses to hits. This effect mirrors the enhanced perceptual sensitivity observed in microstimulation experiments [4, 20].

It is important to note that these results are not capable of resolving the debate about the attentional effects on criterion and sensitivity. Rather, they show that the effects of attention on criterion and sensitivity can be quite complex, and possibly task dependent. For example, the criterion modulation shown in Figure 16 G and H is a result of cue information being transmitted to working memory, where this information has changed the structure of the internal representations in VWM. The cue information has changed the decodability of activated memory transmitted to the reinforcement learning module, affecting behavior in a way that results in a criterion shift. Bias at the time of change mostly results in sensitivity changes. However, bias assignment at the time of change involves multiple interactions. When activated memory patches are drawn from the VWM patches at the time of change, they interact with immediate visual inputs to bias attention on the spatial internal representations associated with the cue location. Moreover, to observe an orientation change there must be stored information of the original stimulus orientations in working memory (since these are drawn randomly). This information also interacts with visual inputs to assign bias to potential change locations. These results and the dynamics displayed by our model demonstrate that criterion and sensitivity effects can involve multiple overlapping pathways and interactions between working memory and attention.

### References

[1] Brian A Anderson. The attention habit: How reward learning shapes attentional selection. *Annals of the new York Academy of Sciences*, 1369(1):24–39, 2016.

[2] Brian A Anderson, Hiroto Kuwabara, Dean F Wong, Emily G Gean, Arman Rahmim, James R Brašić, Noble George, Boris Frolov, Susan M Courtney, and Steven Yantis. The role of dopamine in value-based attentional orienting. *Current Biology*, 26(4):550–555, 2016.

[3] Maximilian Beck, Korbinian Pöppel, Markus Spanring, Andreas Auer, Oleksandra Prudnikova, Michael Kopp, Günter Klambauer, Johannes Brandstetter, and Sepp Hochreiter. xlstm: Extended long short-term memory. *arXiv preprint arXiv:2405.04517*, 2024.

[4] James Cavanaugh, Bryan D Alvarez, and Robert H Wurtz. Enhanced performance with brain stimulation: attentional shift or visual cue? *Journal of Neuroscience*, 26(44):11347–11358, 2006.

[5] Peter Dayan and Nathaniel D Daw. Decision theory, reinforcement learning, and the brain. *Cognitive, Affective, & Behavioral Neuroscience*, 8(4):429–453, 2008.

[6] Robert Desimone, John Duncan, et al. Neural mechanisms of selective visual attention. *Annual review of neuroscience*, 18(1):193–222, 1995.

[7] Michel Failing and Jan Theeuwes. Selection history: How reward modulates selectivity of visual attention. *Psychonomic bulletin & review*, 25(2):514–538, 2018.

[8] Paul W Glimcher. Understanding dopamine and reinforcement learning: the dopamine reward prediction error hypothesis. *Proceedings of the National Academy of Sciences*, 108(supplement_3):15647–15654, 2011.

[9] Priyanka Gupta and Devarajan Sridharan. Presaccadic attention does not facilitate the detection of changes in the visual field. *PLoS Biology*, 22(1):e3002485, 2024.

[10] Okihide Hikosaka, Kae Nakamura, and Hiroyuki Nakahara. Basal ganglia orient eyes to reward. *Journal of neurophysiology*, 95(2):567–584, 2006.

[11] Sepp Hochreiter and Jürgen Schmidhuber. Long short-term memory. *Neural computation*, 9(8):1735–1780, 1997.

[12] Clay B Holroyd and Michael GH Coles. The neural basis of human error processing: reinforcement learning, dopamine, and the error-related negativity. *Psychological review*, 109(4):679, 2002.

[13] Eric I Knudsen. Fundamental components of attention. *Annu. Rev. Neurosci.*, 30(1):57–78, 2007.

[14] Christof Koch and Shimon Ullman. Selecting one among the many: A simple network implementing shifts in selective visual attention. 1984.

[15] Yuan Chang Leong, Angela Radulescu, Reka Daniel, Vivian DeWoskin, and Yael Niv. Dynamic interaction between reinforcement learning and attention in multidimensional environments. *Neuron*, 93(2):451–463, 2017.

[16] Seung-Lark Lim, John P O'Doherty, and Antonio Rangel. The decision value computations in the vmpfc and striatum use a relative value code that is guided by visual attention. *Journal of Neuroscience*, 31(37):13214–13223, 2011.

[17] Thomas Zhihao Luo and John HR Maunsell. Attentional changes in either criterion or sensitivity are associated with robust modulations in lateral prefrontal cortex. *Neuron*, 97(6):1382–1393, 2018.

[18] John HR Maunsell. Neuronal representations of cognitive state: reward or attention? *Trends in cognitive sciences*, 8(6):261–265, 2004.

[19] John HR Maunsell. Neuronal mechanisms of visual attention. *Annual review of vision science*, 1(1):373–391, 2015.

[20] Tirin Moore and Katherine M Armstrong. Selective gating of visual signals by microstimulation of frontal cortex. *Nature*, 421(6921):370–373, 2003.

[21] Yael Niv. Reinforcement learning in the brain. *Journal of Mathematical Psychology*, 53(3):139–154, 2009.

[22] Yoni Pertzov and Masud Husain. The privileged role of location in visual working memory. *Attention, Perception, & Psychophysics*, 76:1914–1924, 2014.

[23] Sebastian Schneegans and Paul M Bays. Neural architecture for feature binding in visual working memory. *Journal of Neuroscience*, 37(14):3913–3925, 2017.

[24] Wolfram Schultz and Anthony Dickinson. Neuronal coding of prediction errors. *Annual review of neuroscience*, 23(1):473–500, 2000.

[25] Richard S Sutton. Reinforcement learning: An introduction. *A Bradford Book*, 2018.

[26] John K Tsotsos. A 'complexity level'analysis of immediate vision. *International journal of computer vision*, 1(4):303–320, 1988.

[27] Freek van Ede, Sammi R Chekroud, and Anna C Nobre. Human gaze tracks the focusing of attention within the internal space of visual working memory. *Journal of Vision*, 19(10):133b–133b, 2019.

[28] Lupeng Wang, James P Herman, and Richard J Krauzlis. Neuronal modulation in the mouse superior colliculus during covert visual selective attention. *Scientific Reports*, 12(1):2482, 2022.

[29] Mary E Wheeler and Anne M Treisman. Binding in short-term visual memory. *Journal of experimental psychology: General*, 131(1):48, 2002.
